# Adaptive translational pausing is a hallmark of the cellular response to severe environmental stress

**DOI:** 10.1101/2020.10.10.334375

**Authors:** Raul Jobava, Yuanhui Mao, Bo-Jhih Guan, Dawid Krokowski, Erica Shu, Di Hu, Evelyn Chukwurah, Jing Wu, Zhaofeng Gao, Leah L. Zagore, William C. Merrick, Youwei Zhang, Xin Qi, Eckhard Jankowsky, Ivan Topisirovic, Donny D. Licatalosi, Shu-Bing Qian, Maria Hatzoglou

## Abstract

Mammalian cells have to adapt to environmental challenges that range from mild to severe stress. While the cellular response to mild stress has been widely studied, how cells respond to severe stress remains unclear. We show here that under severe stress conditions, cells induce a transient hibernation-like mechanism that anticipates recovery. We demonstrate that this <u>A</u>daptive <u>P</u>ausing <u>R</u>esponse (APR) is a coordinated cellular response that limits ATP supply and consumption though mitochondrial fragmentation and widespread pausing of mRNA translation. This pausing is accomplished by ribosome stalling at translation initiation codons, which keeps mRNAs poised to resume translation upon recovery from severe stress. We further show that recovery from severe stress involves adaptive ISR (Integrated Stress Response) signaling that in turn permits cell cycle progression, resumption of growth, and reversal of mitochondria fragmentation. Our findings indicate that cells can respond to severe stress through the APR, a mechanism that preserves vital elements of cellular function under harsh environmental conditions.

## Introduction

The modulation of protein synthesis rates via the regulation of mRNA translation is a necessary and common feature in cellular responses to diverse stress conditions. In addition to playing a major role in shaping the proteome, mRNA translation is also a highly energy-intensive process (Roux and Topisirovic, 2018). To this end, cells adapt to different stressors by disrupting the translational apparatus and altering the translatome to selectively express a subset of factors governing stress-responses (Roux and Topisirovic, 2018). This is in general accompanied by the downregulation of total protein synthesis, which contributes to the maintenance of cellular energy balance under stress. Two major pathways that lead to translational reprogramming in response to a broad range of stresses (e.g. nutrient deprivation, osmotic stress, hypoxia, and ER stress) are the integrated stress response (ISR) and the mechanistic/mammalian target of rapamycin (mTOR) (Costa-Mattioli and Walter, 2020; Roux and Topisirovic, 2018). The ISR involves signaling by eukaryotic translation initiation factor 2 (eIF2) that culminates in the phosphorylation of its α subunit (Costa-Mattioli and Walter, 2020). This impairs recycling of eIF2:GDP to productive eIF2:GTP by the guanine nucleotide exchange factor (GEF) eIF2B, thus limiting Met-tRNAi delivery to the ternary complex (Costa-Mattioli and Walter, 2020). mTOR, a serine/threonine kinase that is a component of two functionally and structurally distinct complexes (mTORC1 and mTORC2) integrates a number of extracellular stimuli and intracellular cues to promote cellular growth and proliferation (Liu and Sabatini, 2020). The mTORC1 complex also plays a major role in regulation of mRNA translation (Roux and Topisirovic, 2018). In response to stress, mTOR signaling is typically downregulated, thereby altering the levels and/or functions of a number of translational and associated factors (Roux and Topisirovic, 2018). For example, decreased mTOR activity is accompanied by activation of a family of small translational repressors, 4E-binding proteins (4EBPs), which impede eIF4F assembly and the m^7^G cap-dependent recruitment of mRNA to the ribosome (Pelletier and Sonenberg, 2019). Inhibition of the mTORC1/S6 kinase axis leads to activation of elongation factor 2 (eEF2) kinase, which phosphorylates eEF2, thus attenuating translation elongation (Leprivier et al., 2013). Stressors, therefore, can induce the ISR and reduce mTORC1 activity to establish adaptive mRNA translational programs, whereby selective translation of downstream target mRNAs is thought to be conferred by specific features of their 5’-UTRs (Hinnebusch et al., 2016). Recent reports have indicated that there are multiple mechanisms that may account for stress-induced perturbations in translation, including, but not limited to phase separation (Franzmann et al., 2018; Iserman et al., 2020), use of alternate initiation factors (Sendoel et al., 2017; Starck et al., 2012), and use of alternate initiation mechanisms (Jeong et al., 2019; Meyer et al., 2015). All the above mechanisms have been widely studied in mild and moderate stress conditions. The molecular strategies that cells use to survive extreme stress are largely unknown. In fact, it is widely believed that harsh environmental conditions kill cells.

The environmental stress of increased extracellular osmolarity (hyperosmotic stress) is a model system to study severe stress (defined by osmolarity concentration and duration). Regulation of cell volume is critical for mammalian life (Burg et al., 2007; Danziger and Zeidel, 2015). In response to increased extracellular osmolarity, cells initially shrink, causing a disruption in mitochondria structure (Copp et al., 2005), a decrease in protein synthesis (Saikia et al., 2012), and alterations in the cytoskeleton (Bustamante et al., 2003). The cellular osmoadaptive response includes induced expression of chaperones and proteins that either synthesize or mediate the cellular uptake of compatible osmolytes, including amino acids (Burg et al., 2007). Therefore, osmoadaptation can remodel the transcriptome and the proteome as a means of accommodating the flux of compatible osmolytes, and restoring cell volume and cellular functions (Grady et al., 2014; Izumi et al., 2015). We have recently shown that increasing stress intensity inhibits osmoadaptation and activates a proinflammatory gene expression program that associates with increased cell death (Farabaugh et al., 2020). Previous studies have suggested that cells sense environmental stress intensity and adjust their responses in order to survive extreme conditions (Saikia et al., 2014). Examples of cells surviving severe stress environments, can be considered in neurodegenerative diseases (Motori et al., 2020), in cancer stem cell survival (Picco et al., 2017) and cancer cell persisters in chemotherapies (Hangauer et al., 2017).

We have here investigated the cellular response to near lethal extracellular osmolarity, a harsh environmental stress condition. We make the striking observation that severe stress induces a hibernation-like cellular state that anticipates recovery from stress. This is a novel cellular response, which we propose to name the <u>A</u>daptive <u>P</u>ausing <u>R</u>esponse (APR). The hallmark of the APR is wide-spread inhibition of mRNA translation initiation, which is accomplished through equally wide-spread ribosome pausing at initiation codons. In addition, we find that cells limit ATP supply through mitochondrial fragmentation, indicating coordination of multiple cellular processes during the APR and adaptation of the cells to prolonged periods of severe stress. We further show that ribosome pausing on initiation codons keeps the cells poised to resume cellular activity during recovery from stress, which we have also characterized. Recovery from the APR exhibits adaptive ISR signaling, ultimately permitting progression of the cell cycle, resumption of growth, and reversal of mitochondria fragmentation. Our findings show how mammalian cells preserve vital elements of cellular function in response to severe environmental stress and reveals the requirement of cellular plasticity to survive harsh environments.

## Results

### Increasing intensity of hyperosmotic stress prevents adaptive recovery of protein synthesis

We examined the proliferation of mouse embryonic fibroblasts (MEFs) in response to increasing hyperosmotic stress conditions (500, 600, and 700 mOsm) over a period of 24 h. In response to 500 mOsm stress, cells adapted and resumed proliferation (Figure 1A). This adaptation to survive in hyperosmotic stress conditions is known as osmoadaptation (Burg et al., 2007). In contrast, in response to 600 and 700 mOsm, osmoadaptation was not observed. Instead, cell proliferation was halted in 600 mOsm, and survival gradually decreased in 700 mOsm stress conditions after 6 h of treatment (Figure 1A). These data indicated that cells can survive severe hyperosmotic stress for a limited time, thus prompting us to investigate the mechanisms of survival in severe environmental stress conditions. We first determined the rates of protein synthesis in MEFs exposed to increasing extracellular osmolarity by [^35^S]-Met/Cys incorporation (Figure S1A). Exposure to mild (500 mOsm) stress for 30 min decreased protein synthesis by 72%, and protein synthesis returned to normal levels with the establishment of osmoadaptation (Figure 1A). Moderate (600 mOsm) and severe (700 mOsm) stress intensities decreased protein synthesis by more than 80% after 2 h, with no osmoadaptive recovery (Figures 1B and S1A). The inhibition of protein synthesis correlated with the increased phosphorylation of eIF2α and dephosphorylation of the mTORC1 substrate p70 (S6K) (Figure 1C). Exposure to 700 mOsm stress as short as 15 min was sufficient to trigger both of these signaling mechanisms (Figure S1B). These data suggest that both eIF2α phosphorylation-mediated decrease in ternary complex availability and mTORC1 inhibition may contribute to the decline of protein synthesis in response to osmotic stress. However, the difference in cellular survival between 500 mOsm and 600 or 700 mOsm (Figure 1A) indicates that cells can proliferate in mild osmotic stress conditions. In contrast, higher stress intensities appear to trigger different mechanisms of delaying decrease in cell survival (Figure 1A, compare 500 mOsm to 600 and 700 mOsm).

**Figure 1.**
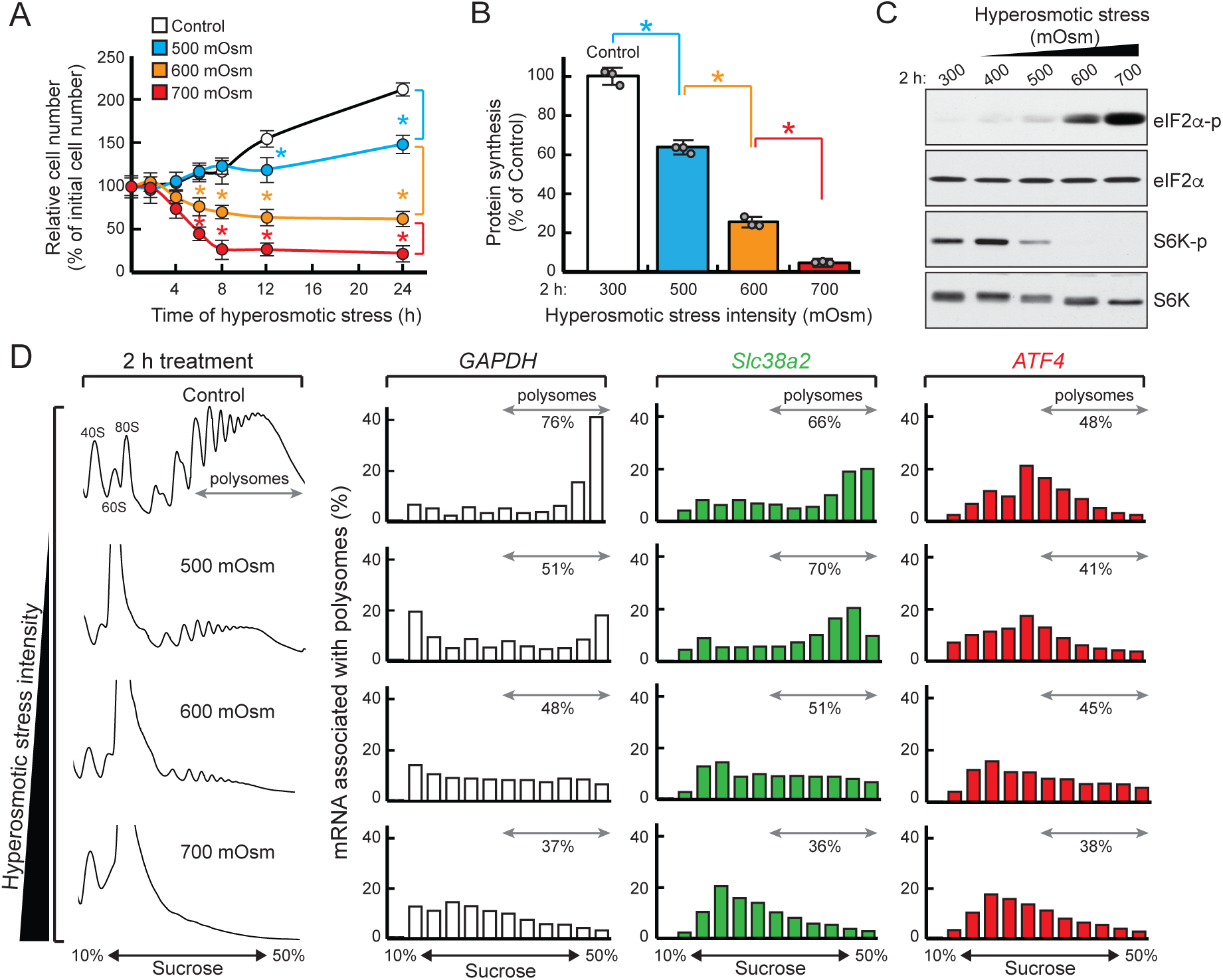
Progressive inhibition of mRNA translation with increasing intensity of hyperosmotic stress. (A) Cell counting in MEFs treated with the indicated hyperosmotic stress conditions. The means ± SEM of triplicate determinations are shown. *p < 0.01 (B) [^35^S]-Met/Cys incorporation into proteins in MEFs treated for 2 h with the indicated hyperosmotic stress conditions. The means ± SEM of triplicate determinations are shown. (C) Western blot analysis for the proteins shown in MEFs treated with the indicated hyperosmotic stress intensities for 2 h. (D) Distribution of the indicated mRNAs on polysome profiles isolated from MEFs treated with the indicated hyperosmotic stress intensities for 2 h.

To compare mechanisms of adaptation in moderate vs. severe stress, we analyzed how global rates of protein synthesis relate to the translation of osmoadaptive and stress-related mRNAs. We monitored the polysomal distribution of mRNAs encoding two master regulators of the cellular response to stress: the osmodaptive amino acid transporter SNAT2/*Slc38a2* (Burg et al., 2007), and the ISR transcription factor ATF4 (Andreev et al., 2015). SNAT2 is a plasma membrane solute carrier which promotes survival under conditions of increased osmolarity via uptake of neutral amino acids (Franchi-Gazzola et al., 2006; Krokowski et al., 2017). Polysome profile analysis revealed the gradual loss of polysomes as stress intensity increased, with concomitant accumulation of monosomes (Figure 1D), consistent with the reduction in global protein synthesis. While some polysomes were preserved in the 600 mOsm condition, 700 mOsm stress caused almost complete dissociation of ribosomes from mRNAs. The house-keeping *GAPDH* mRNA, which we used as a marker of “bulk” translation, shifted to smaller polysomes, 80S, and ribosome-free fractions with increasing stress intensity. In contrast, *Slc38a2* mRNA was translated efficiently in cells exposed to mild stress, but its association with ribosomes progressively declined at more severe stress intensities (Fig 1D). Interestingly, despite high levels of eIF2α phosphorylation, which are known to increase *ATF4* mRNA translation under diverse stress conditions (Andreev et al., 2015), translation of the *ATF4* mRNA was rather attenuated at high intensity of hyperosmotic stress (Figure 1D).

These findings point to a unique translational reprogramming under hyperosmotic stress conditions. We next investigated whether these changes could be reversible by monitoring recovery from 700 mOsm stress after 2 h, at which time cells were reverted to isoosmolar conditions. Hyperosmotic stress is known to induce a DNA damage response that culminates in cell cycle arrest (Dmitrieva and Burg, 2008). We evaluated this aspect of the cellular response to severe hyperosmotic stress using Western blot analysis of proteins involved in the DNA damage response and cell cycle arrest. A replicative stress response (i.e. replication checkpoint) was observed, as evidenced by the significantly elevated phosphorylation of serine/threonine protein kinase checkpoint kinase 1 (Chk1) (Bartek and Lukas, 2003) during early recovery from stress, but not during the stress treatment itself (Chk1-p, Figure S1C). This signal was only activated transiently, as it rapidly decreased after 3 h recovery (Figure S1C). Hence, recovery from hyperosmotic stress immediately activated a DNA replication checkpoint to temporarily inhibit S-phase progression, possibly to allow time for repair of damaged DNA. In contrast, recovery of protein synthesis facilitated the synthesis of p21 and subsequently of cyclin D1 (Figure S1D). Increased p21 levels can induce a G1 and G2/M phase block, whereas the later increase in cyclin D1 can promote progression of cells into S phase and reentry into the cell cycle (Abbas and Dutta, 2009). Notably, during recovery from 2 h of severe hyperosmotic stress, mTORC1 was reactivated as illustrated by increased phosphorylation of p70 (S6K) and 4EBP1 (Figure S1D), which was accompanied by a decline in phosphorylation of eIF2α (Figure S1C). Collectively, these findings suggest that cells halt protein synthesis and growth in response to severe hyperosmotic stress, in anticipation of returning to normal proliferation following removal of the stress. We termed this prosurvival adaptive mechanism to severe stress the <u>A</u>daptive <u>P</u>ausing <u>R</u>esponse (APR).

### Survival in response to increasing stress intensity involves reshaping of the translatome

To better understand the APR, we investigated whether regulation of mRNA translation is involved in restricting cellular function under severe hyperosmotic stress. We performed ribosome profile analysis, which leverages next-generation sequencing to analyze the distribution of ribosome footprints at a sub-codon resolution (Ingolia et al., 2009; Steitz, 1969). We chose to examine ribosome profiling upon exposure of MEFs to hyperosmotic stress of increasing intensity. At 2 h of 500 mOsm stress, we expected cells to be in transition to osmoadaptation (Figure 1A). In contrast, cells after 600 and 700 mOsm stress were expected to be committed to halting cellular activities. Ribosomal footprints and cytoplasmic RNAs were isolated from cells treated with these three different stress intensities, and libraries were prepared and subjected to next-generation sequencing. We evaluated the fold change of RNA levels and ribosome footprints as compared to untreated controls, and classified mRNAs into seven groups in each stress condition (Supplemental File 1): (i) ribosome occupancy (ribo^ocp^) up or (ii) ribosome occupancy down, wherein the ribosome footprint changes exceeded the changes in mRNA levels; (iii) congruent up and (iv) congruent down, exhibiting parallel changes up or down, respectively, in both mRNA levels and ribosome footprints; (v) buffering up or (vi) buffering down, if the changes in mRNA levels exceeded changes in ribosome footprints; and (vii) no change (NC), including the remaining mRNAs which did not exhibit enough change to reach the threshold for the six regulated groups (Figure 2A and Supplemental File 1). This method of analysis can indicate relative differences in mRNA translation within each stress condition, despite the severe inhibition of protein synthesis with increasing stress intensity.

**Figure 2.**
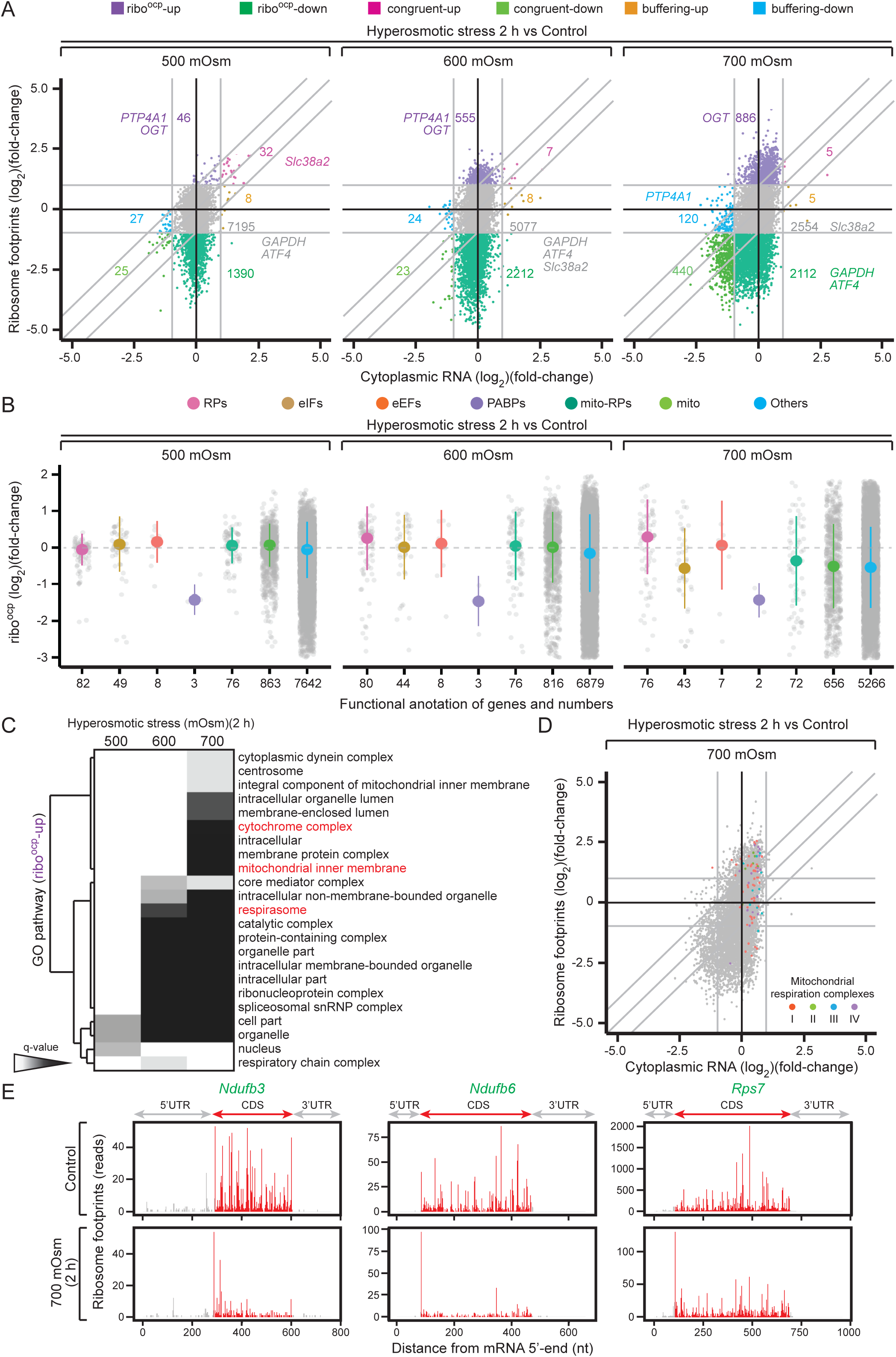
Adaptation to increasing stress intensity involves differential enrichment of mRNAs associated with ribosomes. (A) Scatter plot comparing fold changes of ribosomal footprints (y-axis) and cytoplasmic RNA levels (x-axis) isolated from cells exposed to the indicated hyperosmotic stress intensities for 2 h. The fold change of RNA levels and ribosome footprints as compared to untreated controls were evaluated, and seven groups for each stress condition were generated: (i) ribosome occupancy (ribo^ocp^) up or (ii) down, exhibiting ribosome footprint changes exceeding the changes in the mRNA levels; (iii) congruent up and (iv) down, exhibiting parallel changes up or down, respectively, in mRNA and ribosome footprints; (v) buffering up or (vi) down, exhibiting changes in mRNA levels exceeding the changes in ribosome footprints; and (vii) NC, no change, including the remaining mRNAs which do not exceed the cut-off for the other six regulated groups. Categories are color-coded and the number of mRNAs in each group is indicated. Select mRNAs are identified. (B) Distribution of select groups of mRNAs according to changes in their ribo^ocp^. The number of mRNAs in each group is indicated on the x-axis. (C) Hierarchical clustering of q-values associated with each enriched GO terms (category Cellular Compartment) of the ribo^ocp^-up groups of the indicated stress intensities. Selected GO categories are highlighted in red. (D) Scatterplot as in Figure 2A, comparing changes of the ribosome footprints and cytoplasmic RNA levels, with the mRNAs encoding mitochondrial respiration complexes (I-IV) highlighted. (E) Distribution of ribosome footprints across the mRNA sequences of selected mRNAs under the indicated treatment conditions. Footprints in the coding (CDS) region are highlighted in red and in the untranslated regions (UTRs) in grey.

In agreement with the dramatic decrease in protein synthesis with increasing stress intensity, we observed a gradual loss of mRNAs; 8,723 mRNAs were detected in 500 mOsm stress conditions, 7,906 mRNAs in 600 mOsm, and 6,122 mRNAs in 700 mOsm. The ribo^ocp^-down group in non osmoadaptive stress conditions (600 and 700 mOsm) became larger as the stress intensity increased (Figure 2A). These data suggest that increasing stress intensity leads to decreased association of mRNAs with ribosomes via inhibition of translation initiation of select mRNAs. In agreement with the data in Figure 1D, the mRNAs for *GAPDH* and *ATF4* shifted from the NC group in 500 and 600 mOsm stress conditions to the ribo^ocp^-down group in the 700 mOsm stress condition (Figure 2A). In contrast, the mRNA for *Slc38a2* (Figure 1D) shifted from the congruent up group in 500 mOsm to the NC group in 600 and 700 mOsm stress conditions. These specific examples further demonstrate a hierarchy in selective mRNA translation with increasing stress intensity.

To better understand the nature of selective mRNA translation during adaptive pausing, and since mTOR signaling was severely inhibited by hyperosmotic stress (Figure 1), we examined the translation of mRNAs which have been reported to be positively regulated by mTOR activity; these include mRNAs bearing a 5’-terminal pyrimidine tract (TOP mRNAs), mRNAs demonstrating inhibition of their translation upon treatment with the mTOR competitive inhibitor Torin-1 (Thoreen et al., 2012), and nuclear expressed mRNAs encoding mitochondrial proteins (neMito mRNAs) (Calvo et al., 2016; Morita et al., 2013). We expected to find these mRNAs either in the ribo^ocp^-down group or not bound to ribosomes in high stress intensities. Contrary to our expectations, mTOR signaling-related mRNAs were largely in the NC group in 500 and 600 mOsm stress conditions (Figure S2A and Supplemental File 2). Interestingly, in 700 mOsm, we observed a shift of some mTOR-related and neMito mRNAs to the ribo^ocp^-up group, suggesting that there is additional selection for translation initiation of these mRNAs under severe stress (Figure S2A and Supplemental File 2). We further broke down the mTOR-related mRNAs into more specialized classes: mRNAs coding ribosomal proteins (RPs), eukaryotic initiation factors (eIFs), eukaryotic elongation factors (eEFs), poly(A)-binding proteins (PABPs), nuclear-encoded mitochondrial ribosomal proteins (mitoRPs), and other mitochondrial protein-encoding transcripts. Except for the three PABPs, mTOR-related mRNAs were not present in the low ribosome occupancy groups in 500 and 600 mOsm stress conditions (Figure 2B and Supplemental File 2). These data suggest that a large group of mRNAs maintains translation initiation during severe stress conditions, with a select group of mTOR-related and neMito mRNAs being less inhibited (ribo^ocp^-up) within each stress intensity. The selective mRNA translation of mTOR-sensitive mRNAs was also supported by eIF4E-dependent translation of the mRNA encoding the mitochondrial fission protein MTFP1 (Morita et al., 2017), which was absent in the riboseq analysis at 700 mOsm stress, but present in 500 and 600 mOsm stress conditions (NC group).

Next we more closely examined the ribo^ocp^-up mRNAs. Pathway analysis of the ribo^ocp^-up groups across these three stress intensities (Supplemental File 1) identified mRNAs in the GO category of the “respirasome” enriched in 600 and 700 mOsm but not 500 mOsm stress conditions (Figure 2C). Strikingly, an enrichment of the cellular respiration “cytochrome complex” and “mitochondria inner membrane”-encoding mRNAs were observed in 700 mOsm but not in 500 or 600 mOsm stress conditions, suggesting continuous reprogramming of mRNA translation with increasing stress intensity, to adjust the cellular response to meet shifting needs for survival. This was further supported by KEGG pathway analysis of the ribo^ocp^-up group, which revealed different pathway enrichments in mild versus severe stress condition (Figure S2B).

We next examined the distribution of mRNAs encoding mitochondrial respiration complexes (I-IV) and their assembly factors in the entire translatome in 700 mOsm stress conditions. We identified 85 mRNAs (almost all of the mRNAs in complexes I-III, and half of those in complex IV) distributed mostly in the NC and ribo^ocp^-up groups in the 700 mOsm stress translatome (Figure 2D). However, this distribution only compares relative mRNA translation within the 700 mOsm translatome; the ribo^ocp^-up group indicates better translation than the NC and the ribo^ocp^-down groups, but not better translation than non-stressed cells, in agreement with the severe inhibition of protein synthesis in 700 mOsm stress conditions (Figure 1). We therefore further examined the distribution of footprints within individual mRNAs encoding mitochondrial respiration complexes in the ribo^ocp^-up group. In two representative examples (*Ndufb3* and *Ndufb6* mRNAs), ribosome footprints were distributed in the open reading frames (ORFs) in control cells, indicating efficient translation. In response to 700 mOsm stress, a sharp increase in ribosome pausing at the translation initiation codon was observed, while a lower number of ribosome footprint reads were observed in the body of the ORFs (Figure 2E). mTOR-related mRNAs from the NC group, such as *rpS7*, also showed a dramatic decrease in the distribution of footprints in the ORF and pausing at the translation initiation codon (Figure 2E). Taken together, these data suggest that adaptive pausing during severe hyperosmotic stress may halt ribosomes at the translation initiation codons of a select group of mRNAs. Ribosome pausing may also serve as a mechanism for swift recovery in the event that the environmental stress is removed.

### Transitioning to adaptive pausing is marked by changes in the translatome and mRNA abundance of a select group of mRNAs

The pathway analysis of the ribo^ocp^-up mRNAs in different stress intensities (Figure 2C) suggested remodeling of the mRNA translation selectivity with stress intensity. Of the mRNAs in the ribo^ocp^-up group at 500 mOsm, 57% remained in the same category at 600 mOsm, and of the mRNAs in the ribo^ocp^-up group at 600 mOsm, 74% remained at 700 mOsm. In cells exposed to 500 mOsm stress for 2 h, the relative number of ribosome footprints increased in 46 mRNAs without concomitant upregulation of their mRNA levels (ribo^ocp^-up, Figure 2A). Among the mRNAs with the highest changes in ribosome occupancy were *PTP4A1* (Protein tyrosine phosphatase type IV A 1) which regulates the TGF-β signaling pathway (Sacchetti et al., 2017), known to be important for hyperosmotic stress signaling (Tew et al., 2011), and *OGT* (O-Linked N-Acetylglucosamine (GlcNAc) Transferase), which post-translationally modifies proteins involved in the stress response (Groves et al., 2013) (Figures 2A and S2C). We confirmed the translational upregulation of these mRNAs at 500 mOsm by RT-qPCR analysis of the individual fractions in the polysome profiles (Figure S2C). Both mRNAs remained associated with ribosomes in higher stress intensities, in support of their known functions in the stress response (Groves et al., 2013; Hardy et al., 2019).

In addition to translational control, in 500 mOsm stress conditions, 32 mRNAs were in the congruent up group, which had significant increases in both mRNA levels and ribosome footprints (Figure 2A). KEGG pathway analysis revealed the enrichment of terms such as ‘TNF signaling’, ‘MAPK signaling’, and ‘IL-17 signaling’ (Figure S2B). Examination of the individual genes that made up these pathways suggested that negative feedback regulators of the p38 kinase and NF-κB signaling pathways (*DUSP1*, *NFKBIA*, *TNFAIP3*, and *IER3*) were among the κ top congruent up mRNAs, which is in agreement with the termination of inflammation as the cells progress to an osmoadaptive state (Farabaugh et al., 2020). These data suggest that progression to osmoadaptation involves changes in mRNA abundance and translational control to promote expression of osmoadaptive genes and inhibit signaling that promotes stress sensing and inflammatory gene expression programs. In contrast, stress-induced adaptive pausing revealed “ribosome” and “oxidative phosphorylation” as the predominant KEGG pathways, which include enrichment of ribosomal proteins and nuclear-encoded mitochondrial proteins (Figure S2B). Finally, APR at 700 mOsm was marked by 440 mRNAs in the congruent down group (Figure 2A). These mRNAs were primarily in the NC and ribo^ocp^-down mRNA groups in 500 and 600 mOsm stress conditions. Taken together, these data suggest that increasing stress intensity applies additional layers of translational and mRNA abundance control, engaging the limited cellular resources for translation of those mRNAs in greater need, in anticipation of recovery from the environmental stress condition. Translation initiation codon pausing (Figure 2E) is likely an additional layer of regulation of selective mRNA translation during APR. This prompted us to further investigate the patterns of the distribution of ribosome footprints on individual codons throughout the entire translatome in 700 mOsm stress conditions.

### Ribosome stalling at translation initiation codons of specific mRNAs during severe hyperosmotic stress

To obtain a genome-wide view of ribosome pausing at translation initiation codons with increasing stress intensity, we evaluated the average ribosome density around the authentic translation initiation AUG and around translation termination codons (Figure 3A). This analysis showed elevated ribosome density at the initiation AUG codon, but not at the termination codon, when cells are subjected to severe hyperosmotic stress. It should be noted that the occupancy at the ribosomal P-site was used for analysis, as ribosome occupancy was not changed at the A-site. This result suggests that the codon decoding process is not likely affected by stress, but that the ribosome movement is halted when an AUG codon is at the ribosome P-site. These data also indicate that severe stress promotes translation initiation of a select group of mRNAs, but 80S ribosome pausing at initiation codons will halt productive synthesis of the corresponding proteins. Analysis of ribosome occupancy on individual codons showed that, on average, ribosomes spend more time at the initiation codon in isoosmolar conditions, in agreement with this being a slow step in the transition from translation initiation to elongation (Wang et al., 2019). In agreement with the analysis in Figure 3A, pausing at AUG initiation codons increased with stress intensity (Figure 3B). Pausing at other codons across all ORFs was similar between isoosmolar and stress conditions. Interestingly, we also observed that 60% of the mRNAs with paused ribosomes contained footprints at AUG codons (and non-cognate initiation codons) downstream of the authentic translation initiation codons (Figure 3C). It is well known that in stress conditions characterized by decreased ternary complex availability, authentic translation initiation codons can be skipped due to leaky scanning of the 40S ribosome (Andreev et al., 2015). Since the footprints marked P-site AUG pausing, we therefore hypothesize that the observed pausing events within the main ORFs may be the result of failed translation initiation events at the authentic initiation codons due to leaky scanning (Andreev et al., 2015).

**Figure 3.**
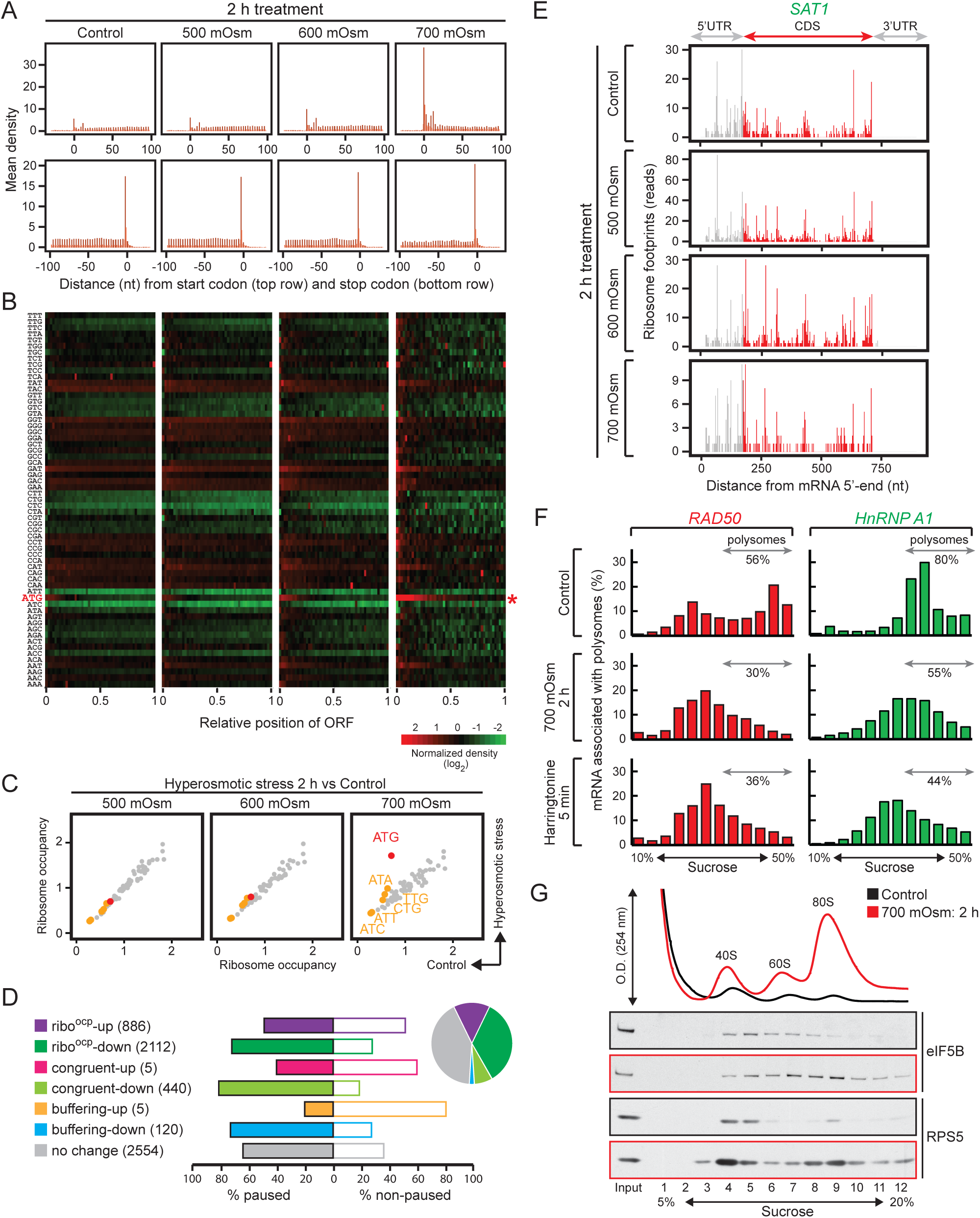
Sustained inhibition of protein synthesis during severe hyperosmotic stress is accompanied by pausing of ribosomes at AUG translation initiation codons of a select group of mRNAs. (A) Mean density of ribosomal footprints relative to the translation initiation AUG codon (top row) and the translation termination codon (bottom row) in MEFs exposed to the indicated stress conditions. (B) Heat map depicting ribosome occupancies on the different codons (left) based on their relative location on the mRNAs (x-axis). The AUG codon is highlighted in red. (C) Scatter plots comparing ribosome occupancies of the different codons in ORFs, excluding authentic translation initiation codons, between isoosmolar and the indicated stress conditions. Putative translation initiation codons within the ORFs are highlighted in red (AUG) and green (near cognate CUG, UUG, GUG) colors. (D) Percentage of paused and non-paused mRNAs in each group of mRNAs identified in cells treated with 700 mOsm for 2 h (Figure 2A). The groups of mRNAs are color-coded, and the number of mRNAs in each group is indicated and represented in a pie-chart. (E) Distribution of footprints on *SAT1* mRNA in cells exposed to the indicated hyperosmotic stress conditions. Footprints in the CDS are highlighted in red, and in the UTRs in grey. (F) Distribution of individual mRNAs (*RAD50* and *hnRNPA1*) on polysome profiles of cytoplasmic extracts isolated from MEFs treated with the indicated stress conditions. The data are representative of three biological replicates. (G) Western blot analysis for the indicated proteins of sucrose gradient fractions from cells treated with the indicated stress conditions and cross-linked in 1% formaldehyde. The positions of the 40, 60, and 80S ribosomes are marked. Data are representative of three independent experiments.

In order to compare the relative pausing between different mRNA groups engaged with ribosomes in severe stress conditions, we calculated a pausing index for individual mRNAs by dividing the number of ribosome reads at the translation initiation AUG codons (combined authentic and downstream AUG initiation codons) by the reads in the remainder of the ORF (Supplemental File 3). Paused mRNAs (pausing index ≥ 2) were identified among each of the 7 mRNA groups (Figure 3D), suggesting that in response to severe 700 mOsm hyperosmotic stress, ribosome pausing at translation initiation codons marks a suspension of translation initiation. The lower enrichment in paused mRNAs in the ribo^ocp^-up group (Figure 3D) is in agreement with better translation of a select group of mRNAs during severe stress. The data in Figure 2E serve as an example of ribo^ocp^-up paused mRNAs at 700 mOsm stress. A striking example of a non-paused mRNA in the ribo^ocp^-up group at all stress intensities (Figure 2A) was the polyamine metabolism spermidine/spermine N1-acetyl transferase 1, *sat1* (Brett-Morris et al., 2014) (Figure 3E). Translation of *sat1* mRNA increased in 500 mOsm stress conditions, and some translating ribosomes were maintained in the main ORF without strong pausing at the translation initiation codon even in 700 mOsm stress conditions (compare Figs. 3E and 2E). We further identified the enrichment of genes of the “ribosome” and “oxidative phosphorylation” pathways in the 700 mOsm ribo^ocp^-up group in both paused and non-paused mRNAs (Figure S3A). These data suggest that the degree of pausing differs in the pool of mRNAs translated during severe stress.

We hypothesized that paused mRNAs with very low translation (ribo^ocp^-down group) will be enriched on the 80S ribosomes on polysome profile gradients in 700 mOsm stress conditions. We further hypothesized that this distribution would be similar to the effects of the translation initiation inhibitor harringtonine on non-stressed cells. Harringtonine inhibits protein synthesis by allowing runoff of polyribosomes and freezing initiation at monosomes (Ingolia et al., 2011). As an example, we focused on the mRNA *RAD50* encoding a DNA repair protein (Dmitrieva and Burg, 2008), which is in the NC group in 500 and 600 mOsm stress conditions (Figure 2A). Compared to isoosmolar conditions, during 700 mOsm *RAD50* mRNA showed an enrichment in fractions 5-7, which correspond to 80S ribosomes (Figure 3F). The distribution of *RAD50* mRNA was indistinguishable between harringtonine-treated samples and those after 700 mOsm hperosmotic stress (Figure 3F). In contrast to *RAD50*, the paused mRNA for *HnRNPA1* in the NC group at 700 mOsm was associated with ribosomes heavier than 80S during stress, suggesting some translation of this mRNA was maintained during severe stress (Figure 3F). These data suggest that translation initiation of select mRNAs is halted during severe stress, most likely due to the slow first step in translation elongation.

To identify the cause of ribosome pausing at the translation initiation AUG codon, we first determined if aminoacylation of the initiator Met-tRNA (Met-tRNA_i_) was affected by severe hyperosmotic stress. No change in aminoacylation of the Met-tRNA_i_ was observed in any of the three stress intensities tested (Figure S3B), suggesting that there is no limitation of the Met-tRNA_i_ for translation initiation at AUG codons. We next investigated a recently reported rate-limiting step in translation initiation, release of the translation initiation factor eIF5B (Wang et al., 2019). eIF5B plays an important role in translation initiation by mediating 40S and 60S subunit binding, as well as stabilizing Met-tRNA_i_ binding at the P site of the ribosome (Shin et al., 2002). In some systems, eIF5B was also shown to deliver the Met-tRNA_i_ to the P site of the ribosome (Wang et al., 2019). After 80S assembly on the AUG initiation codon at the P site, and before the first elongator tRNA is delivered to the A site, translation initiation factor eIF5B must be released (Lee et al., 2002). eIF5B has intrinsic GTPase activity which is stimulated by the 60S GTPase center; this hydrolysis event is necessary for the dissociation of eIF5B from the 80S ribosome (Shin et al., 2002). Release of eIF5B from the ribosome is a rate-limiting step in translation initiation (Olsen et al., 2003; Wang et al., 2019). We therefore tested if eIF5B is present in the paused 80S ribosome fractions in polysome profiles from cells in the 700 mOsm stress condition. In isoosmolar conditions, eIF5B migrated in the same fractions as the 40S ribosomes, while at 700 mOsm, eIF5B shifted to the fraction of the 80S ribosomes. These data suggest that ribosome stalling at translation initiation codons during severe hyperosmotic stress may involve a slowed release of eIF5B from the initiating ribosome (Figure 3G).

### Stress duration-dependent checkpoints during severe hyperosmotic stress dictate the kinetics of recovery

As shown earlier, cells reverted to isoosmolar media after 2 h treatment with 700 mOsm stress media can re-enter the cell cycle (Figure S1D). In order to understand the significance of ribosome pausing after cells exit the adaptive pausing state, we evaluated whether the return of cells to isoosmolar conditions coincided with reversal of ribosome pausing at translation initiation sites. Indeed, mRNAs whose translation was paused upon 700 mOsm stress (as per Figure 3D and Supplemental File 3) associated with more ribosomes during the first 15 min and up to 2 h of recovery from stress (Figure 4A). To further confirm that the degree of pausing on mRNAs during stress may reflect their efficiency in initiating translation during recovery, we compared the top 20% of mRNAs in which ribosomes were paused at authentic AUG initiation codons (highest pausing index) vs. the bottom 20% (lowest pausing index) of the same group of mRNAs (Supplemental File 4). We found that mRNAs with a higher degree of pausing were also more preferentially translated upon recovery (Figure S4A). This was further shown by the distribution of ribosome footprints in the mitochondrial respiration complex mRNAs *Uqcrsf1* (top 20%) and *Ndufv1* (bottom 20%). Ribosome pausing at the translation initiation AUG codon recovered after 2 h in isoosomolar media, with ribosome footprints in these mRNAs returning to control levels (Figure S4B). This was further confirmed with the return of the ribo^ocp^ values of respirasome-associated mRNAs to the levels of untreated cells (Figure 4B). These data indicate that (i) ribosome pausing occurs on mRNAs exhibiting translation initiation during severe stress, and (ii) paused mRNAs have a small advantage in being translated ahead of non-paused mRNAs during recovery from stress.

**Figure 4.**
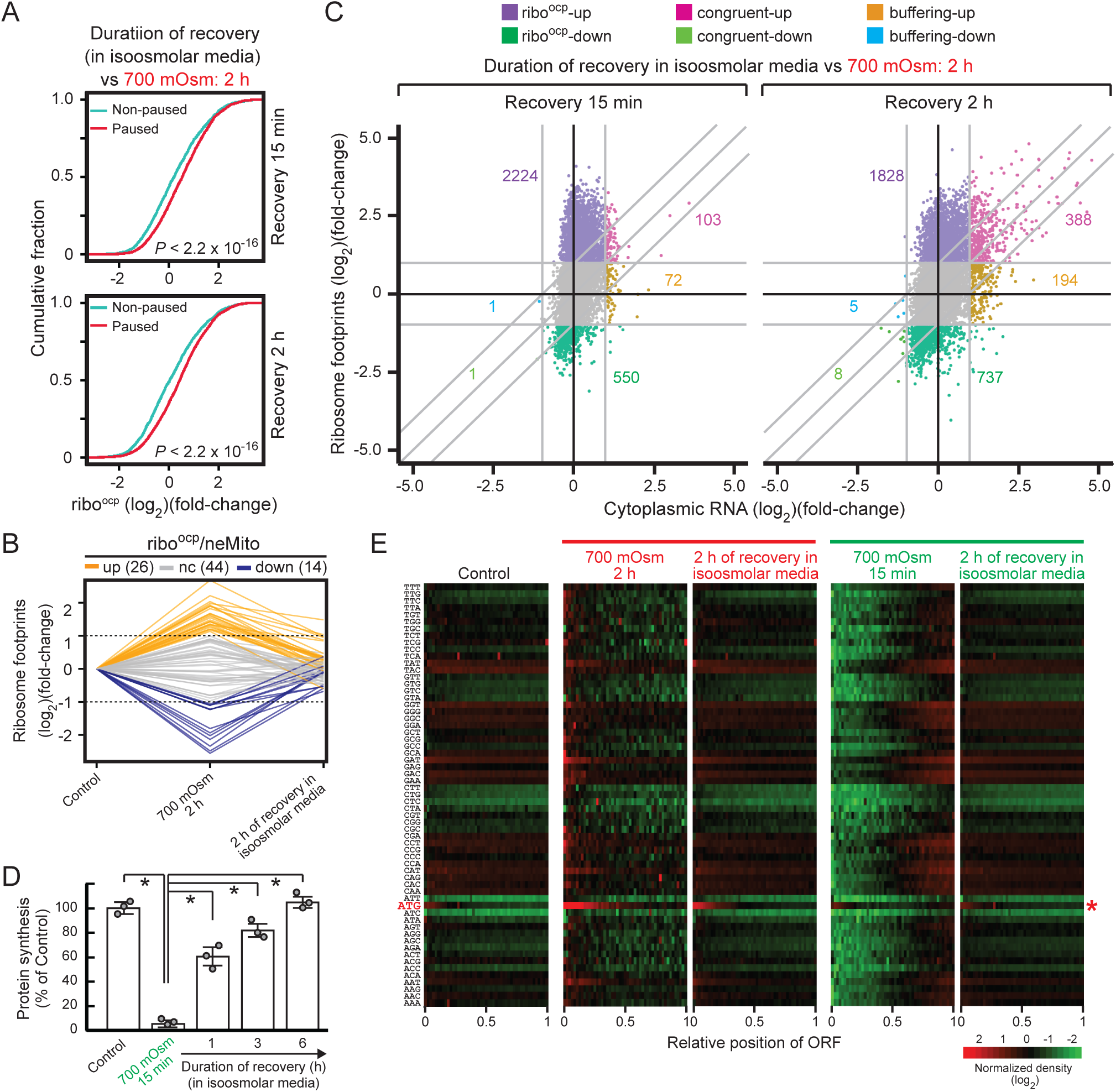
Pausing at translation initiation AUG codons is stress duration-dependent and reversed by the return of cells to isoosmolar conditions. (A) The cumulative distribution function (y-axis) of the fold change of ribo^ocp^ values (x-axis) between stress (700 mOsm, 2 h) and recovery (15 min (top) and 2 h (bottom)) was calculated for mRNAs that were either paused (red) or non-paused (cyan) in 700 mOsm 2 h (Figure 3D). (B) Relative changes of the ribo^ocp^ values of the neMito mRNAs that encode genes of the respirasome, in transition from isoosmolar to 700 mOsm (2 h) stress and recovery to isoosmolar conditions (2 h). Changes in three groups are indicated: ribo^ocp^-up (up), no change (nc), and ribo^ocp^-down (down). (C) Scatter plot comparing fold changes of ribosomal footprints (y-axis) and cytoplasmic RNA levels (x-axis) of the cells after recovering for 15 min or 2 h in isoosmolar media from 700 mOsm stress (2 h). The comparisons were made to the values of 700 mOsm stress (2 h). Groups of mRNAs are color-coded, and the number of mRNAs in each group is indicated as in Figure 2A. (D) ^35^S-Met/Cys incorporation into proteins in MEFs treated with 700 mOsm (15 min) and recovered in isoosmolar media for the indicated durations. The means ± SEM or triplicate determinations are shown. *p < 0.01. (E) Heat map depicting ribosome occupancies on the different codons (y-axis) based on their relative location on mRNAs (x-axis). The AUG codon is highlighted in red.

To further understand the dynamics of gene regulation during the first 2 h of recovery from hyperosmotic stress, we analysed the trancriptomes and translatomes of the recovery from 700 mOsm stress after 15 min and 2 h (Figure 4C). During recovery, the distribution of the ribo^ocp^ values suggested massive reversal of the protein synthesis inhibition observed after 2 h of 700 mOsm stress (compare Figs. 2A and 4C). However, the opposite was also observed; of the 550 mRNAs with decreased ribo^ocp^ during recovery (Figure 4C), 85% were in the ribo^ocp^-up group after 2 h of 700 mOsm stress. Reprogramming of mRNA abundance was evident in the congruent up group (103 mRNAs), which was enriched in genes of the ‘transcription’ and ‘cellular signaling’ pathways (Figure S4C). Interestingly, 93% of these mRNAs were in the congruent down group in 700 mOsm stress conditions (compare Figs 2A and 4C). A similar observation was made for the buffering up group (72 mRNAs, Figure 4C), which were primarily in the congruent and buffering down groups in 700 mOsm stress conditions. These data suggest that in the first 15 min after exiting adaptive pausing, cells reverse the translational and mRNA abundance reprogramming in response to severe stress, while a transcription and signaling program develops to promote a return to homeostasis. This conclusion was further supported by the significant increase of mRNAs in the congruent up and buffering up groups after 2 h of recovery from adaptive pausing (Figure 4C, 2 h recovery). Interestingly, 396 mRNAs in these groups were in the ribo^ocp^-up group after 15 min of recovery. Taken together, these data suggest that the early response in recovery from adaptive pausing is focused on regulation of mRNA translation, while the late response involves congruent changes in the early response translatomes.

This concept of time-dependent programs that function upon the exit of adaptive pausing has been previously described in the adaptation of cells to chronic ER stress(Guan et al., 2017). However, in contrast to adaptation to chronic ER stress, in which cells adapt to survive under the stress condition, recovery from adaptive pausing is a transient state prior to the return to homeostasis. We therefore put forth two hypotheses: (i) adaptive pausing develops in a stress duration-dependent maner, and (ii) exiting the adaptive pausing state likely shares similar elements with the cellular response of adaptation to chronic ER stress. To test the first hypothesis, cells were exposed to 700 mOsm for 15 min, which inhibited protein synthesis by 95%, similar to the inhibition shown after 2 h of 700 mOsm stress (compare Figure 1B and Figure 4D). The subcodon resolution of ribosome distributions showed clearance of ribosomes at the 5’-end of the ORFs and accumulation near the stop codons (Figure 4E). Since the average elongation rate in mammalian cells is 5-6 codons/sec (Ingolia et al., 2011; Wu et al., 2016), even ORFs as long as 6 kb should have been devoid of ribosomes by 6-7 min, suggesting that severe hyperosmotic stress decreases translation elongation (Burg et al., 2007) in addition to inhibition of translation initiation. The most prominent group of mRNAs with ribosomes in the 3’-halves of the ORFs were those mRNAs with ORFs longer than 1 kb (Supplemental File 1). This is clearly demonstrated by the mRNAs for fibronectin (*FN1*) and *Rad50* (Figure S4D). Similar to our earlier observations (Figure 3D), the global pausing at translation initiation codons after 2 h of stress, was not evident after 15 min of stress (Figure 4E). These data suggest that the development of ribosome pausing on translation initiation codons in response to severe hyperosmotic stress is stress duration-dependent. Furthermore, these data suggest that there is a temporal component to the development of selective mRNA translation initiation during severe stress, with ribosome pausing at translation initiation codons comprising an additional layer of translational control. Overall, the data also suggest that APR is a stress intensity- and duration-dependent cellular response to severere hyperosmotic stress. We next compared the dynamics of gene regulation in response to 15 min stress and recovery. Hyperosmotic stress for 15 min induced dramatic translational reprogramming (Figure S4E and Supplemental File 1). The overall patterns observed in the transcriptome/translatome of cells recovering from this short duration of 700 mOsm stress were similar to the cells recovering from exposure to the longer duration of 2 h of stress (Figure S4E). Translational recovery of 2047 mRNAs during early recovery was followed by increased mRNA abundance (69 mRNAs congruent up and 69 buffering up). However, both, the number of mRNAs and the magnitude of increased abundance were smaller in cells recovering after 15 min stress compared to the cells recovering from 2 h of stress (compare Figures 4C and S4E). These data further suggest that cells sense the duration of exposure to severe stress, and adjust the magnitude of recovery accordingly. This conclusion converges on our second hypothesis, that recovery from long durations of severe hyperosmotic stress has elements similar as the adaptation to chronic ER stress (Guan et al., 2017). Both conditions have duration-dependent development of adaptation programs; the early response involves translational control, while the late response involves a new transcriptome which was in part derived from the early response-regulated translatome.

### Cellular sensing of the duration of severe hyperosmotic stress induces stress adaptation programs promoting a return to homeostasis

As shown above (Figure 4A-E), cells sense stress duration and intensity, and adjust their transcriptomes and translatomes in anticipation of recovery from stress and the return to homeostasis. It is expected that this return to homeostasis and resumption of growth would be swift during recovery from short durations of severe stress, and slower following adaptive pausing. Therefore we hypothesized that recovery following adaptive pausing involves induction of stress adaptation mechanisms prior to resumption of cell growth. To test this hypothesis, we compared changes in the transcriptomes and translatomes between control and recovering cells in the following two conditions: (i) 700 mOsm stress for 15 min followed by recovery for 2 h and 700 mOsm stress for 2 h followed with recovery for 2 h, compared to control isoosmolar conditions (Figure 5A). Upon recovery from short stress, cells showed nearly complete return to homeostasis, with 36 mRNAs in the ribo^ocp^ down group, and 50 mRNAs in the combined congruent up and ribo^ocp^-up groups (Figure 5A). In contrast, after recovery from long stress duration, cells maintained the stress-response patterns of ribo^ocp^-down (927 mRNAs), ribo^ocp^-up (185 mRNAs), and congruent up (186 mRNAs). In order to further understand the effect of duration of stress on the return of cells to homeostasis upon recovery, we compared the functional significance of mRNAs that were more associated with ribosomes (combined ribo^ocp^ and congruent up groups) during 2 h of recovery from either short or long stress durations. The 50 mRNAs that had higher ribosome association in the recovery period after short stress duration showed enrichment of genes involved in “Cytokine-cytokine receptor interaction”, “IL-17 signaling pathway”, “TNF signaling pathway”, and the “MAPK signaling pathway” (Fig 5B). This is in agreement with proinflammatory signaling promoting growth, as previously described in other cell injury systems (Werner and Grose, 2003). Furthermore, the presence of pro-growth signaling was evident from the association of proliferation mRNA *Ereg* (Singh et al., 2016), mitochondria oxidative capacity mRNA *Angptl4* (Guo et al., 2014), and growth factor mRNA *VEGF* (Apte et al., 2019) with the highest fold change of ribosome occupancy (Figure 5A). The 371 mRNAs that displayed higher association with ribosomes after recovery from 2 h of 700 mOsm stress conditions showed enrichment in “Transcriptional misregulation in cancer”, “MAPK signaling pathway”, and “TNF and FoxO signaling pathways” (Figure 5B). Interestingly, a large number of transcription factors were encoded by these mRNAs (Figure 5A), suggesting that during recovery from prolonged stress, the synthesis of transcription factors facilitates reprogramming of gene expression for resumption of growth and survival. A prominent ISR program was evident, as *ATF4*, *ATF5*, *ATF3*, and *CHOP* were among these 371 mRNAs, in addition to downstream targets such as the cystine transporter *SLC7A11* (Koppula et al., 2018) and the E3 ligase *HERPUD1* (Li et al., 2018). In agreement with a regulated transition from adaptive pausing to homeostasis is the presence among the 371 mRNAs of mRNAs associated with the termination of inflammatory signaling (*NFKBIA* and *TNFAIP3*) and dual-specificity phosphatases (DUSPs 1, 2, 4, 6, 8, 10, 14 and 16) with known functions in the termination of ERK/MAPK signaling and inhibition of apoptosis (Lang and Raffi, 2019). Therefore cells exiting APR, enter a prosurvival-stress adaptation program that combines transcriptional and translational control mechanisms of regulation of gene expression.

**Figure 5.**
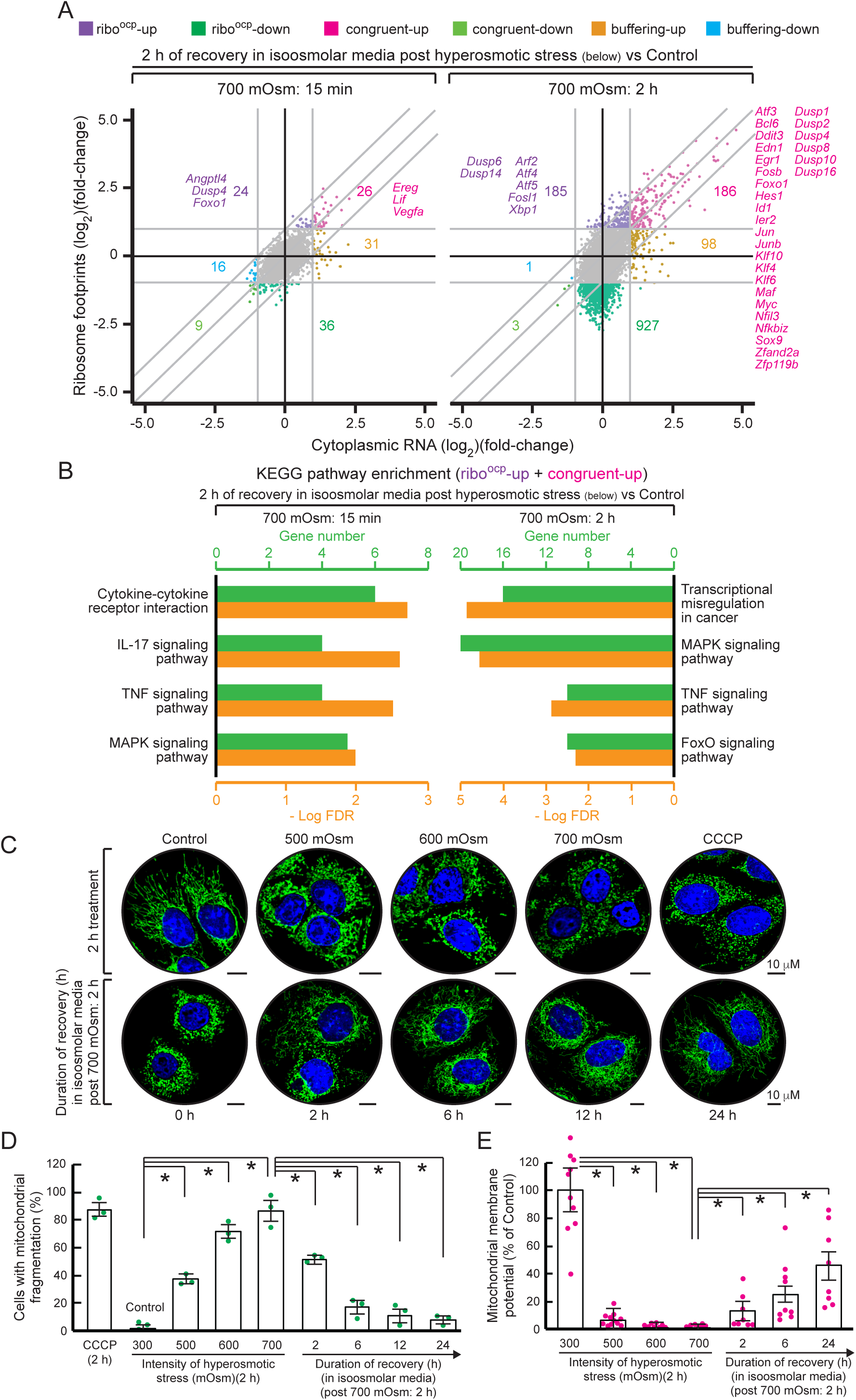
Re-shaping of the translatome/transcriptome in recovery from severe stress is determined by the stress duration. (A) Scatter plot comparing fold changes of ribosomal footprints (y-axis) and cytoplasmic RNA levels (x-axis) of the cells recovering in isoosmotic media (2 h) from 700 mOsm (15 min (left) or 2 h (right)). The comparisons were made relative to control (isoosmotic) conditions. The mRNAs belonging to different categories are highlighted, and the number of mRNAs in each group is indicated. Selected mRNAs important for the stress response are indicated and color-coded. (B) KEGG enrichment pathway analysis of mRNAs with increased ribosome association (ribo^ocp^-up + congruent up). (C) Representative confocal microscopy images of mitochondria (Tom20, green) and the nucleus (Hoechst, blue) in MEFs treated as indicated. Treatment for 2 h with the inhibitor of oxidative phosphorylation, CCCP, served as a positive control for mitochondrial fragmentation. Scale bar: 10 μm. (D) Quantification of cells with fragmented mitochondria from confocal microscopy images treated as indicated. The means ± SEM of triplicate determinations are shown. *p < 0.01. (E) Mitochondrial membrane potential determined in MEFs treated as indicated. The means ± SEM of triplicate determinations are shown. *p < 0.01.

The enrichment in paused mRNAs encoding mitochondrial protein subunits of the respiration complexes (Figs. 2D and 4B) prompted us to examine mitochondrial morphology in a stress intensity-dependent manner and during recovery from APR. Exposure of MEFs to increasing stress intensity caused mitochondrial fragmentation, with 40% of cells having fragmented mitochondria in 500 mOsm stress and almost 90% at 700 mOsm stress conditions (Figure 5C-D). Mitochondrial uncoupler carbonyl cyanide chlorophenylhydrazone (CCCP) as a postive control (Lou et al., 2007) induced mitochondrial fragmentation in about 90% of MEFs, comparable to the extent of fragmentation observed at at 700 mOsm stress (Figure 5D). When cells were reverted to isoosmolar media after 2 h of 700 mOsm stress, mitochondrial fragmentation decreased to 50% after 2 h of recovery, and the tubular mitochondrial network almost completely recovered after 24 h (Figs. 5C-D). Consistent with this, we observed a stress intensity-dependent decrease in mitochondrial membrane potential (MMP) in MEFs, whereas a gradual increase in the MMP was observed during recovery from stress (Figure 5E). The severe decrease in MMP (Figure 5E) further supports that the effort of the cells in severe stress intensities to maintain functionality by initiating translation of a select group of mRNAs is halted by a negative-feedback mechanism of inhibition of productive protein synthesis. Taken together, these data suggest that translational pausing during severe hyperosmotic stress impacts mitochondrial morphology and function.

### Slow kinetics of recovery from severe hyperosmotic stress reveal the induction of stress-adaptation mechanisms

We noted the induction of an ISR gene expression program during recovery from adaptive pausing (Figure 5A). It is well known that induction of the ISR requires a decrease in eIF2B activity, leading to a reduced level of ternary initiation complex availability; this in turns inhibits global protein synthesis and activates translation of a select group of mRNAs, among them the transcription factor ATF4 (Han et al., 2013). The ISR is a prosurvival mechanism of adaptation to diverse stress conditions (Guan et al., 2017). The presence and function of the ISR in recovery from severe hyperosmotic stress are largely unexplored. We first investigated protein synthesis rates when cells were reverted from 700 mOsm stress to isoosmolar media (Figure 6A). Incorporation of [^35^S]-Met/Cys into proteins was gradually restored, reaching approximately 40% by 6 h of recovery and almost 100% by 12 h of recovery (Figure S5A). The slow kinetics of recovery of protein synthesis (Figure 6A) were not consistent with the reactivation of mTOR, as shown by the phosphorylation of p70 (S6K) (Figure 6B). In contrast, the gradual recovery of global protein synthesis correlated precisely with the gradual recovery of eIF2B activity (Figure 6C). eIF2B activity, which was severely inhibited upon stress, remained inhibited for the first 2 h of the recovery period, and only began returning to normal levels thereafter (Figure 6C). We also determined if the delayed recovery of eIF2B activity induced the adaptive ISR response (Costa-Mattioli and Walter, 2020; Guan et al., 2017). Although the ISR was not active during hyperosmotic stress (Figure 1), despite low eIF2B activity (Figure 6C), it was activated during recovery from severe stress (Figure 5A). Induction of both ATF4 and GADD34 increased significantly (Figure 6B) upon recovery. GADD34 is a subunit of the PP1 phosphatase, which is involved in dephosphorylation of eIF2α and recovery of stress-induced inhibition of protein synthesis in all adaptive stress-induced cellular responses (Novoa et al., 2001). In agreement with our previous studies (Guan et al., 2017), very low levels of eIF2α phosphorylation were sufficient to sustain a low eIF2B activity during stress recovery, and thus, induce the ISR (Figure 6B). Polysome profile analysis confirmed the increased association of *ATF4* mRNA with polysomes after 2 h of recovery from stress (Figure 6D). Taken together, the recovery from severe hyperosmotic stress demonstrates similar elements to the cellular response to stressors that promote adaptation to diverse stress conditions (Pakos-Zebrucka et al., 2016), including ER stress (Guan et al., 2017), via the well-known mechanism of ISR (Costa-Mattioli and Walter, 2020). These data further suggest that severe hyperosmotic stress induces a pausing cellular state which suspends stress-induced adaptive mechanisms, despite the critical signaling of decreased eIF2B activity. In contrast, stress-induced adaptive mechanisms are transiently activated during recovery from pausing, thus introducing the importance of the ISR in this process.

**Figure 6.**
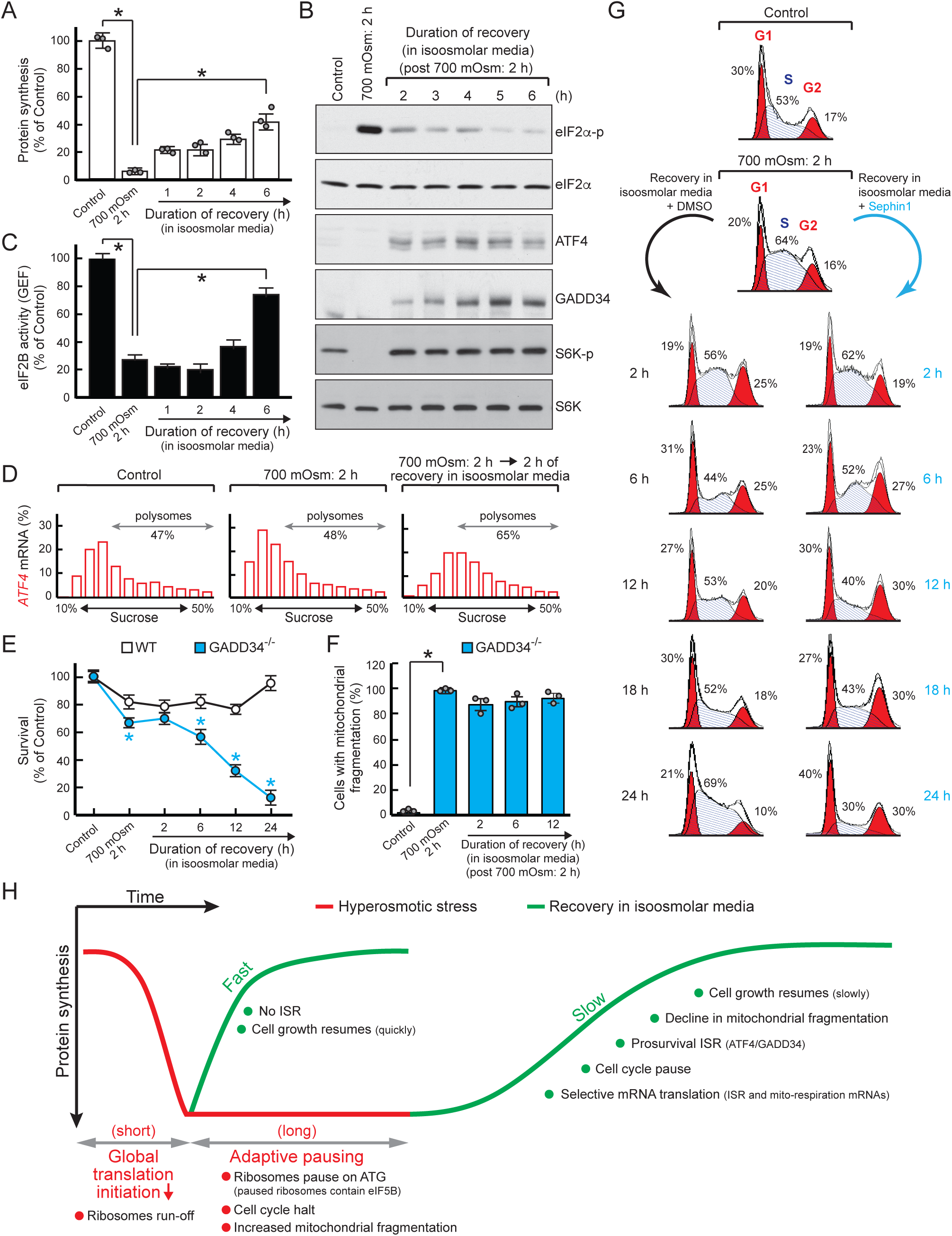
ISR is a hallmark of the recovery from adaptive pausing in response to severe stress. (A) [^35^S]-Met/Cys incorporation into proteins in MEFs treated with the indicated stress conditions. The means ± SEM or triplicate determinations are shown. *p < 0.01. (B) Western blot analysis of the time course of recovery from 700 mOsm stress (2 h). Data are representative of three independent experiments. (C) eIF2B GEF activity of cellular extracts of the MEFs exposed to the indicated treatments. The means ± SEM or triplicate determinations are shown. *p < 0.01. (D) ATF4 mRNA distribution on polysome profiles of cells treated with the indicated stress conditions. The percentage of mRNA distributions in the indicated fractions is shown. (E) Cell survial measured for the indicated cells and the indicated times with Cell-titer glo kit (Promega). The means ± SEM or triplicate determinations are shown. *p < 0.01. (F) Quantification of cells with fragmented mitochondria from confocal microscopy images treated as indicated. The means of ± SEM of triplicate determinations are shown. *p < 0.01. (G) Flow-cytometric analysis of propidium-iodide stained cells in the presence or absence of GADD34 inhibitor sephin1. Cell cycle phases (and percentage of cells in each phase) are indicated. (H) Temporal cellular response to severe hyperosmotic stress. Severe stress for short durations is characterized by global ribosome run-off, followed by swift recovery upon return to isoosmotic conditions. During prolonged stress, cells enter the “ribosome pausing” state characterized by translation initiation and pausing of a select pool of mRNAs, followed by slow recovery of cellular functions upon return to isoosmotic media. Recovery from prolonged stress is accompanied by cell cycle arrest and the induction of the ISR. Reversible mitochondrial fragmentation is a hallmark of the cellular response to prolonged hyperosmotic stress.

### Induction of the ISR-related GADD34/PP1 during recovery from severe hyperosmotic stress promotes cell survival

As several APR characteristics are not observed after only short exposures to severe hyperosmotic stress, we investigated whether the ISR is also absent during recovery from short durations of severe hyperosmotic stress. We first observed that recovery of protein synthesis was swift after 15 min of stress (Figure 4D), in contrast to the slow recovery of protein synthesis after 2 h of severe stress. Similarly, we did not observe expression of genes of the ISR program (Figure S5B). In agreement with this faster isoosmolar recovery of protein synthesis, we observed accumulation of the short-lived CCAAT Enhancer Binding Protein (CEBP) LIP (Li β et al., 2008). The reinstated CEBP β LIP level marks the recovery of protein synthesis in isoosmolar conditions after 15 min of hyperosmotic stress. These data suggest that the entry and exit from the adaptive pausing state have unique elements that prepare cells to delay apoptosis and recover once the stress resides. To address this hypothesis, we analyzed the well-known prosurvival axis of ISR, the function of GADD34 in the recovery from severe hyperosmotic stress.

The best-characterized function of GADD34 is its ability as a regulatory subunit of the PP1 phosphatase to dephosphorylate eIF2α, and thus alleviate a stress-induced inhibition of protein synthesis (Brush et al., 2003; Reid et al., 2016). However, functions of GADD34 independent of eIF2α dephosphorylation have also been described, such as in protein trafficking (Krokowski et al., 2017), and in the inhibition of apoptosis (Reid et al., 2016). We therefore investigated whether GADD34-mediates the recovery of protein synthesis after the return of cells to isoosmolar conditions following severe hyperosmotic stress, via the use of GADD34^-/-^ cells (Harding et al., 2009). We first showed that GADD34^-/-^ cells had a similar response with WT cells in the regulation of protein synthesis with increasing stress intensity (compares Figure S5C and 1B). In addition, in GADD34^-/-^ cells, regulation of mTOR-substrate protein phosphorylation in stress and recovery was similar to WT cells (compare Figures 6B and S5D). During hyperosmotic stress recovery, deficiency of GADD34 inhibited restoration of protein synthesis, in agreement with sustained high levels of eIF2α phosphorylation (Figures S5D and S5E). These data suggest that the induction of GADD34 during recovery from severe stress promotes the gradual recovery of protein synthesis via the modulation of the levels of eIF2α phosphorylation. Therefore, the function of ISR in part, is to cause gradual recovery of protein synthesis upon exit from adaptive pausing.

To assess the prosurvival function of ISR during recovery from stress, we determined the impact of GADD34 depletion on cell survival during recovery from severe hyperosmotic stress. In agreement with the inhibited recovery of protein synthesis, GADD34^-/-^ cells had decreased survival during recovery from stress (Figure 6E) and mitochondria fragmentation stayed high (Figure 6F). These data suggest that GADD34/PP1 is required for cell survival and cell cycle re-entry during recovery from severe stress. Therefore we determined the progression of the cell cycle in the presence of sephin1 (Das et al., 2015), an inhibitor of PP1/GADD34-phosphatase activity (Figure 6G). We have previously shown that sephin1 can decrease osmoadaptation via the GADD34/PP1 axis (Krokowski et al., 2017). Thus we used the same inhibitor to study recovery from adaptive pausing. We made the following observations: (i) hyperosmotic stress for 2 h induced a moderate shift of cells from the G1 and S phases to the G2/M phase of the cell cycle, which persisted for the first 2 h of the recovery period; (ii) the 2 h recovery was followed by resumption of the G1 phase, but the reduction in cells in the S phase persisted until 6 h of recovery. Accordingly, the number of cells in the G2/M phase remained high during this period; (iii) after recovery for 6 h, the cells returned to normal cell cycle progression as the number of cells in the S phase increased. (iv) The presence of sephin1 did not prevent the reduction in the number of cells in the S phase from 2-6 h of the recovery period; (v) in the presence of sephin1, cells were not able to return to a normal cell cycle progression, and instead maintained the inhibition entering the S phase. We conclude that inhibition of GADD34/PP1 caused cell cycle arrest in the G1 phase (Figure 6G). Although it is unknown how GADD34 promotes S phase entry, it is likely via its function in recovery of protein synthesis inhibition and amino acid transporter trafficking (Krokowski et al., 2017), as amino acid uptake is required for cell cycle progression (Cano-Crespo et al., 2019). We propose that the induction of ISR during recovery from severe hyperosmotic stress promotes the return of cells to normal growth and survival via the functions that involve the GADD34/PP1. The similar states of the cell cycle at 2 h of 700 mOsm stress and after 2 h of recovery further support that the induction of the ISR and other gene expression programs shown in Figure 5A is due to a recovery from the adaptive pausing response, and not cell cycle-mediated regulation of gene expression.

We conclude that cells sense the duration and intensity of stress, and adjust the efficiency of translation of select mRNAs. In response to prolonged severe stress, cells enter an adaptive pausing state of translational inactivity that impacts mitochondrial function, while maintaining mechanisms to resume growth in anticipation of removal of the stress. During recovery from the adaptive pausing state, cells induce stress-adaptation transcriptional programs and signaling events which reprogram the cellular mRNA content and terminate stress signaling events, allowing re-entry into the cell cycle and resumption of growth. We therefore conclude that adaptive pausing prevents stress-induced apoptosis and preserves the ability of the cells to recover from severe stress (Figure 6H).

## Discussion

In response to hyperosmotic stress, cells either undergo cell cycle arrest followed by adaptation (Arsenijevic et al., 2013) or undergo apoptosis (Farabaugh et al., 2020). The mechanisms by which cells choose between these alternative pathways in response to different stress intensities remains obscure. We show here that cells exposed to severe hyperosmotic stress induce a duration-dependent hibernation-like inhibition of cellular processes, which delays cell death and prepares cells in anticipation of recovery from the stress insult. We have named this transient pro-survival response to severe stress the Adaptive Pausing Response (APR). The APR is characterized by the induction of signaling that inhibits mRNA translation initiation, and promotes severe mitochondria fragmentation. This cytosol-mitochondria communication in severe stress was further supported by the selective translation initiation of a subset of mRNAs enriched in the pathways of protein synthesis and mitochondria function. Despite translation initiation, ribosome pausing at initiation AUG codons was identified as a unique feature of APR. These APR features were reversed when stress was removed and cells returned to isoosmolar conditions. Cells developed the adaptive ISR response during recovery from stress, which allowed cell cycle progression and resumption of growth. We propose that, as showcased here for severe hyperosmotic stress, severe environmental stress induces translational silencing, yet preserves vital elements of cellular function via distinct layers of regulation of mRNA translation. Furthermore, this report provides an experimental system to study the long-standing questions concerning the factors that orchestrate co-regulation of mitochondria energy production, protein synthesis, and proliferation (Morita et al., 2015).

Mitochondria fragmentation indicates a disruption in the balance between fission and fusion processes, which continuously adjust to accommodate cellular energy demands (Liesa and Shirihai, 2013). Mitochondria fragmentation increases in response to cellular stress, and can be adaptive or can induce cell death (Youle and van der Bliek, 2012). We show here that osmotic stress induces reversible mitochondria fragmentation with minimal cell death. The extent of mitochondria fragmentation appeared to be proportional to stress intensity, which was mirrored by the dramatic translational reprogramming of nuclear-encoded mitochondrial genes. Similar to hyperosmotic stress, mitochondria fragmentation in adaptive conditions has been reported in other contexts; for example, mitochondria fragmentation accelerated wound healing in response to tissue injury (Fu et al., 2020), and was essential for metabolic adaptation in hypothalamic neurons for the control of systemic glucose homeostasis (Toda et al., 2016). One potential mechanism by which mitochondria fragmentation can induce an adaptive response of cells to harmful stimuli is by inhibiting inter-organelle Ca^2+^ transport (Szabadkai et al., 2004), thus altering Ca^2+^ signaling (Fu et al., 2020). In addition to the translational mechanisms presented here, it has been proposed that mechanical force-mediated Ca^2+^ signals in response to cell shrinkage induced by hyperosmotic stress can be a potential mechanism for mitochondria fragmentation in osmoadaptation (Kim et al., 2015). The high percentage of mitochondria fragmentation in APR, which was paralleled by a dramatic decrease in the mitochondrial membrane potential and translational pausing of nuclear-encoded mitochondria mRNAs, may be similar to the recently reported requirements of mitochondria fission and low metabolic activity as a mechanism to preserve cellular integrity and self-renewal of hematopoietic stem cells (Hinge et al., 2020; Liang et al., 2020).

The discovery of adaptive pausing revealed that cells use different mechanisms to adapt to stress, in a manner dependent on the exposure time and intensity of the stress. Although it is remarkable that in mild hyperosmotic stress intensity cells reprogram their transcriptomes and translatomes to maintain homeostasis and continue dividing and functioning, it is even more astonishing that cells have the plasticity to continue reshaping their translatomes as the stress intensity increases with the expectation to recover when the stress environment changes. We showed here that translational control was a major mechanism in achieving this plasticity. With varying stress intensities, we observed differential enrichment of cellular pathways in the mRNAs that remained associated with ribosomes, as well as a hierarchy in the efficiency of translation initiation and ribosome pausing of these mRNAs during severe stress. The enrichment and persistent association of mRNAs which are involved in oxidative phosphorylation and protein synthesis with ribosomes during severe stress is in agreement with the idea of maintaining a low-energy metabolic state in preparation for a swift recovery, as was shown here to be the case. It was interesting that of the 21 nuclear-encoded mRNAs for oxidative phosphorylation complex subunits, only one, the complex I subunit *NDUFV2* mRNA, had low ribosome occupancy in stress conditions, indicating lower efficiency for translation initiation compared to the rest of the subunits. Complex I is the entry point for the mitochondrial respiratory electron transport chain (ETC) for ATP synthesis. It is also a major source of ROS production in the ETC, which can be harmful to cells, especially under stress conditions with weak anti-oxidant defense mechanisms (Snezhkina et al., 2019). Although speculative, the low translation efficiency of *NDUFV2* relative to other subunits may indicate a defense mechanism against superoxide radical production. This speculation is supported by the finding that NDUFV2 is expressed in lower levels in longer-lived animals, in association with lower mitochondrial ROS synthesis and reduced symptoms of aging (Mota-Martorell et al., 2020).

The characterization of APR during severe hyperosmotic stress revealed unique elements, as compared to elements of the severe environmental stress of heat shock, which are conserved across species (Richter et al., 2010). In contrast to the translation initiation pausing observed in APR, in response to severe but not lethal heat shock (44° C for 2 h), a translation elongation block was observed within the first 60 nucleotides of most, but not all, mRNAs (Shalgi et al., 2013). This translational pausing was caused by decreased activity and levels of chaperone proteins, in agreement with increased nascent protein ubiquitination during heat shock (Aprile-Garcia et al., 2019). In the heat shock response, translation of the mRNAs encoding ISR transcription factors ATF4 and ATF5 escaped translation elongation pausing, and were translated along with other transcription factors important for the response to severe heat shock (Shalgi et al., 2013). In contrast, in APR, *ATF4* and *ATF5* were translationally repressed, and the reprogramming of transcription was evident during recovery from stress. Another remarkable example of adaptive ribosome stalling was recently described for the stresses of nutrient deprivation and UV radiation. It was shown that the pro-survival ISR is induced via activated protein kinases SAPK and GCN2, via mechanisms that involve colliding ribosomes (Wu et al., 2020).These differences suggest that the APR is a unique adaptive cellular state of severe environmental stress, reminiscent more of hibernation than an active adaptive stress response program to maintain cellular functions under severe stress. Similar to hibernation, APR uses reversible suppression of mitochondrial metabolism to survive extreme environmental stress conditions (Mathers et al., 2017; Staples, 2014).

One noted feature of the APR was the hierarchy of translational efficiency and translation initiation codon pausing among the mRNAs under conditions of severe inhibition of protein synthesis. The latter associated with decreased mTOR activity and increased eIF2α phosphorylation, both inhibitory mechanisms of mRNA translation. Surprisingly, among the mRNAs with higher ribosome occupancy were mRNAs known to be severely inhibited by the inhibition of mTOR (Thoreen et al., 2012). These data suggested that the hierarchy in translation initiation may reflect a hierarchy in eIF4F-mediated mRNA translation. This argument is supported by recent literature indicating that there are at least eight different eIF4F complexes involved in translation initiation (Robert et al., 2020). Our findings indicating a hierarchy in mRNA translation initiation during severe inhibition of protein synthesis in APR are also supported by an elegant report in yeast, in which the authors showed using eIF4E-tagged capture and RNA-seq that acute oxidative stress or nutrient limitation revealed subpopulations of mRNAs with a hierarchy of translation initiation and differential eIF4F depletion. The pool that was translated more efficiently exhibited more complete depletion of eIF4F, when the more translationally repressed group exhibited stabilization of eIF4F protein (Costello et al., 2017). These previous reports are consistent with modeled systems demonstrating mRNA competition for translation initiation (Godefroy-Colburn and Thach, 1981). Taken together, these reports support a model of differential stability of eIF4F complexes with mRNAs in response to acute inhibition of protein synthesis by stress conditions (Costello et al., 2017). Our findings may be explained by the differential stability of eIF4F complexes with mRNAs during APR. It remains for future studies to determine the signaling pathways that can lead to differential ribosome recruitment by APR-translated mRNAs. Towards that end, the function of RNA binding proteins in stress-mediated phase separation (Riback et al., 2017) or hyperosmotic stress-induced cytoplasmic localization of RNA binding proteins (Bevilacqua et al., 2010; Hock et al., 2018) may also be considered as potential mechanisms for APR.

The initiation of the ISR during recovery from the APR had unique features in comparison to cellular adaptation to chronic ER stress (Guan et al., 2017). Similar to adaptation to chronic ER stress, ATF4 was translationally upregulated when eIF2B activity was low, and in turn induced expression of target proteins CHOP and GADD34. In contrast to the chronic ER stress which associates with sustained low eIF2B activity (Guan et al., 2017), the recovery of the hyperosmotic stress-induced inhibition of protein synthesis was dependent on the recovery of the eIF2B activity via dephosphorylation of eIF2α. In this manner, the recovery from the APR is similar to the recovery of cells from transient ER stress, in which the eIF2α kinase is transiently activated and therefore causes the transient decrease of eIF2B activity. The requirement of GADD34 for recovery from APR may also involve mechanisms that we have shown previously for progression into osmoadaptation. It was shown that the GADD34/PP1 activity is required for plasma membrane protein trafficking and osmoprotective protein localization (Krokowski et al., 2017).

The translational control mechanism described here in response to severe hyperosmotic stress may be applicable in other physiological extreme environmental stress conditions. Hibernation in mammals represents one example of adaptation to severe stress. Remarkably, in mammalian hibernation, neurons lose their ability to generate long term potential (LTP), but they maintain an arousal state throughout the hibernation cycle (Horowitz and Horwitz, 2019). Similar observations were recently made for mouse neurons that regulate the response to hypothermic and nutrient restricted environments during torpor (Hrvatin et al., 2020). The latter identified an adaptive energy-conserving prosurvival response, in which a subpopulation of neurons remain responsive to permit termination of torpor. These observations suggest that neuroplasticity is essential for surviving extreme stress conditions. Interestingly, both hibernation and torpor associate with increased phosphorylation of eIF2α (Frerichs et al., 1998) and inactivation of eEF2 (Yamada et al., 2019).

The findings in this study provide insights into the cellular response to extremely harsh environments, and reveal an unexpected response preserving the ability to recover once that environment changes. In addition to the resemblance of the identified APR to the physiological process of hibernation and torpor, another broad significance of these finding lies in diseases such as cancer. Therapeutic strategies to kill cancer cells aim to induce cell death in actively dividing cells with unusually high nutrient demands and deregulated ROS levels (Truitt et al., 2015). However, in response to chemotherapies, cancer cells can enter cellular dormancy, a state resistant to such therapies (Hangauer et al., 2017). Cancer cell dormancy is a reversible state of metabolic inactivity, and can contribute to cancer recurrence and metastasis. Molecular signatures of dormant cells include inactive mTOR signaling (Kim et al., 2013), activation of the p38 kinase pathway and phosphorylation of eIF2α (Ranganathan et al., 2006), and hypoxia-induced signaling (Butturini et al., 2019). Interestingly, the cellular response to hypoxia mimics the hibernation response (Stone et al., 2019). Although the APR is evident in the response to the severe environmental stress of increased extracellular osmolarity, it likely plays a role in other physiological cellular responses and the development of disease.

## Acknowledgements

The authors thank all Hatzoglou lab and Qian lab members for constructive discussions and technical assistance with the experiments. The authors thank the Genomics Core facilities at Cornell Institute of Biotechnology, the Genomics Core at CWRU School of Medicine, and Novogene Co., Ltd, for performing the next-generation sequencing. The authors thank the Cytometry & Imaging Microscopy Shared Resource of the Case Comprehensive Cancer Center and especially Mike Sramkoski for help with the flow-cytometry analysis. This body of work was supported by the following grants: NIH R01DK053307, R01DK060596, and R01DK113196 to MH; CDDRCC pilot grant DK097948 to MH; NIGMS R01GM107331 to DDL; National Science Centre (Poland) 2018/30/E/NZ1/00605 to DK; R21CA227917, R01GM1222814, and HHMI Faculty Scholar (55108556) to S.-B.Q; and the Canadian Institutes for Health Research [MOP-363027] and Joint Canada-Israel Health Research Program (JCIHRP) [108589-001] to I.T., who is a Research Scholar (Senior) of the Fonds de Recherche du Québec—Santé (FRQ-S); YZ supported in part by R01CA230453; D.H. and X.Q. are supported by R01AG065240 and R01NS115903. WCM. is supported by GM128981-0A1.

## Author Contributions

Project conceptualization, R.J. and M.H.; Methodology Development, R.J.,Y.M and B.-J.G..; Investigation, R.J., Y.M., B.-J.G, E.C., J.W., D.H., E.S., L.Z., D.D.L; Bioinformatics, Y.M.; Resources, M.H. and S.-B.Q, ; Data Curation, R.J., Y.M., D.D.L and B.-J.G; Writing, R.J. and M.H.; Review & Editing, R.J., M.H., I.T., D.L., D.K., W.M., E.J., S.-B.Q., X.Q.; Visualization, R.J., B.-J.G. and Y.M.; Funding Acquisition, I.T., S.-B.Q., M.H.; Supervision, S.-B.Q. and M.H.

## Methods

### Experimental Model and Subject Details

#### Cell Lines, Cell Culture Conditions, and Chemicals

The MEFs used were described previously (Scheuner et al., 2001). The cells were grown in high glucose Dulbecco’s modified Eagle’s medium (DMEM) (ThermoFisher Scientific, 11960044) supplemented with 10% fetal bovine serum (ThermoFisher Scientific, 26140079), 1X Penicillin-Streptomycin-Glutamine (ThermoFisher Scientific, 10378016) at 37°C and 5% CO_2_. For all experiments, cells were subcultured 24 hours prior to the experiment so that cells were ∼80% confluent at the time of the experiment. GADD34^-/-^ were a gift from D. Ron (Novoa et al., 2003). Hyperosmotic stress was induced with Sorbitol (Sigma-Aldrich, S1876). Other chemicals used in this study include cyclohexamide (CHX) (100 μg/mL) (Sigma-Aldrich C7698), harringtonine (2 μg/mL) (LKT Laboratories, H0169), sephin 1 (Apexbio,A8708) and CCCP (Sigma-Aldrich, C2759).

#### Protein Extraction and Western Blot Analysis

Cells were washed twice with ice-cold phosphate-buffer saline (PBS) and scraped into lysis buffer (50 mM Tris-HCl at pH 7.5, 150 mM NaCl, 2 mM EDTA, 1% NP-40, 0.1% SDS, and 0.5% sodium deoxycholate) supplemented with EDTA-free protease inhibitor (Roche Applied Science) and PhosSTOP phosphatase inhibitor (Roche Applied Science) (1 tablet of each per 10 mL)), and sonicated briefly 10 times (2 RMS). After centrifugation (12000 X g, 10 min, 4°C), proteins were quantified using the DC Protein Assay Kit (Bio-Rad). Equal amounts of protein 10-20 μg) were analyzed by Western blotting. Primary antibodies used in this study are listed in Key Resources Table.

#### Measurement of protein synthesis rates via metabolic labeling with ^35^S-Met/Cys

Cells on 24-well plates were treated with appropriate stress conditions, followed by incubation with ^35^S-Met/Cys (30 mCi/mL EXPRESS Protein Labeling Mix, PerkinElmer) for an additional 30 min. After quickly washing with ice-cold PBS twice, cells were incubated with 5% trichloroacetic acid (TCA)/1 mM cold methionine for 10 min on ice (repeated three times) and lysed in 200 µL of 1 N NaOH/0.5% sodium deoxycholate for 1 h. Incorporation of radioactive amino acids was determined by liquid scintillation counting and normalized to the cellular protein content (quantified with DC Protein Assay (Bio-Rad)).

#### Flow cytometric analysis of cell cycle via propidium iodide staining of DNA

After treatment, cells were trypsinized, pelleted at 850 X g, and washed with PBS. Cells were fixed by incubation with 70% ethanol overnight at 4°C. The cells were treated with 100 µg/ml RNase A (ThermoFisher Scientific, EN0531). Cells were stained with 50 μg/ml propidium iodide (Sigma, P4170). Staining was analyzed on a FACSAria™ II instrument. ModFit LT was used to fit a model to the data.

#### CellTiter-Glo and proliferation assays

Cell viability was measured with CellTiter-Glo Luminescent Cell Viability Assay (Promega) following the manufacturer’s instructions. For the cell viability assay performed in Figure1A, 5 X 10^5^ cells were seeded on 6 cm^2^ dishes. 24 h post-seeding, cells were treated with 500, 600, or 700 mOsm hyperosmotic media and imaged at 2, 4, 6, 8, 12, and 24 h after treatment. The number of cells still attached to the plate in each field of view at 20X magnification were determined manually from at least 3 images with over 250 cells total. This number of cells was expressed as the percentage of cells counted at the same specifications immediately prior to treatment.

#### Analysis of tRNA charging by gel electrophoresis

After respective treatments, RNA was isolated with Trizol according to the manufacturer’s instructions, except that RNA was dissolved in 10 mM sodium acetate (pH 5)/1 mM EDTA. To prepare deacylated tRNAs, RNA was incubated in 0.2 M Tris-HCl (pH 9.5) for 1 h at 37°C, followed by precipitation with ethanol and resuspension in 10 mM sodium acetate (pH 5)/1 mM EDTA. 10 µg RNA was mixed with equal volume acid urea sample buffer (0.1 M sodium acetate (pH 5), 8 M urea, 0.05% bromophenol blue, 0.05% xylene cyanol FF). RNA was resolved by an acid urea polyacrylamide gel (12% polyacrylamide (19:1 acrylamide/bisacrylamide), 0.1 M sodium acetate (pH 5), 8 M urea) at 500 V over 24 h at 4°C in 0.1 M sodium acetate (pH 5) buffer. GENIE® blotter (Idea Scientific, 4003) was used to transfer to GeneScreen Plus Hybridization Transfer Membrane (PerkinElmer, NEF986001PK) for 30 min in 1X TBE buffer (0.089 M Tris Base, 0.089 M Boric Acid, 0.002 M EDTA). The membrane was UV crosslinked (120 mjoules/cm^2^) using a Stratalinker Spectroline™ XL-1500 (ThermoFisher,11-992-90). Probe was labeled with T4 PNK (NEB M0201S) at 37°C for 1 h in a reaction containing 20 pmol probe, 1X T4 PNK Reaction Buffer, 50 pmol [γ-32P]-ATP (7000 Ci/mmol, 150 mCi/ml)(PerkinElmer, NEG002Z250UC). For hybridization, ULTRAhyb™-Oligo(ThermoFisher Scientific, AM8663) was used according to the manufacturer’s instructions.

#### Measuring *in vitro* guanine nucleotide exchange factor (GEF) activity of eIF2B

GEF activity of the cellular extracts was assayed as previously described (Kimball et al., 1989; Guan et al., 2017). Briefly, after washing twice with ice-cold PBS, cells were lysed in the homogenization buffer (45 mM HEPES-KOH at pH 7.4, 0.375 mM MgOAc, 75 μM EDTA, 95 mM KOAc, 10% glycerol, 1 mM DTT, 2.5 mg/mL digitonin, protease and phosphatase inhibitors (Roche Applied Science; 1 tablet per 10 mL)). Cell lysates were incubated on ice for 15 min, vortexed occasionally, passed six times through a 26-gauge needle, and centrifuged at 13,000 rpm for 15 min at 4°C. After determining protein concentration using DC Protein Assay (Bio-Rad), 100 μg was used for the assay. eIF2 (purified from rabbit reticulocyte lysate) was incubated with ^3^H-GDP for 10 min at 30°C in buffer (62.5 mM MOPS at pH 7.4, 125 mM KCl, 1.25 mM DTT, 0.25 mg/mL bovine serum albumin (BSA)). The complex was stabilized by the addition of 2.5 mM MgOAc and incubated with nonradioactive GDP (0.1 mg/mL)-supplemented lysate at 30°C. At 0, 2, 4, and 6 min, aliquots of the mixture were removed and filtered through nitrocellulose filters. The amount of eIF2α-^3^H-GDP complex bound to the nitrocellulose filters was assessed by liquid scintillation counting. The eIF2B activity was calculated as the rate of exchange of ^3^H-GDP for nonradioactive GDP.

#### Ribosome footprinting analysis

Three 15 cm plates of MEFs (80% confluency) were treated with the indicated intensity of hyperosmotic stress. Cells were washed three times with ice-cold PBS and scraped in 1 mL lysis buffer (10 mM HEPES pH 7.4, 5 mM MgCl_2,_100 mM KCl,1% Triton-X,100 μg/mL cycloheximide, 2 mM DTT, RNase inhibitor (1:100 dilution, NEB, M0314L), protease inhibitor (1 tablet per 10 mL, Roche, 04693159001). The lysate was passed four times through a 26 gauge needle and centrifuged at 1300 x g for 10 min. Lysate (OD A_260_ = 10) was loaded on a linear 10%-50% sucrose gradient (gradient buffer was the same as the lysis buffer, lacking only Triton-X and Protease and RNase inhibitors) and centrifuged at 4°C at 31,000 rpm for 2 h and 30 min using an SW 32 Ti rotor(Beckman Coulter, 369694). The gradients were fractionated using a Teledyne ISCO Density Gradient Fractionation System and 12 fractions were collected. The fractions from (including) the 80S peak to the bottom of the gradient were combined. 300 μL of the combined fractions was digested with 5 μL RNase I (Thermo Fisher Scientific, AM2295) for 1 h at 4°C (slow rotation on the nutator). Immediately, Trizol LS (Thermo Fisher Scientific, 10296028) was added, and RNA isolated according to the manufacturer’s instructions.

10 μg of total RNA isolated from the combined fractions was resolved by 15% TBE-Urea gel (Thermo Fisher Scientific, EC6885BOX) and visualized with SYBR™ Gold Nucleic Acid Gel Stain(Thermo Fisher Scientific, S11494); 28-30 nt region was excised from the gel. The RNA was eluted in 400 μL RNA elution buffer (EB) at 4°C overnigt (*RNA EB*: 0.3M sodium acetate (NaOAc, pH 5.5), 1 mM EDTA, 0.4 U/μL RNase inhibitor). Gel debris was removed using Corning® Costar® Spin-X® centrifuge tube filters (Sigma, CLS8160). RNA was precipitated with 4 µL GlycoBlue™ Coprecipitant (Thermo Fisher scientific, AM9515), 40 µL sodium acetate, and 1 mL ethanol overnight at -80°C. Recovered footprints were dephosphorylated in a 15 µL reaction mixture (1 µL T4 PNK (NEB, M0201S), 10X T4 PNK buffer, 0.4 U/μL RNase inhibitor) at 37°C for 1 h followed by heat inactivation at 65°C for 10 min. Dephosphorylated RNA was precipitated as described above and recovered in 4 µl of water. Pre-adenylated DNA linker(DNA oligo purchase from IDT was adenylated using 5′ DNA Adenylation Kit (NEB,E2610S)) was ligated to the 3’-end of RNA in a 20 µl reaction (25% PEG8000, 1X T4 RNA ligase buffer, 1 mM DTT, 0.4 U/μL murine RNase inhibitor, 300 U T4 RNA ligase truncated KQ (M0373S, NEB), 1 µM pre-adenylated DNA linker) for 6 h at 25°C followed by overnight ethanol precipitation as described above. Ligated footprints were resolved by 10% TBE-Urea Gel (Thermo Fisher Scientific, EC6875BOX), purified, eluted in the RNA EB at 4°C overnight, precipitated at -80°C as described above and resuspended in 10.5 µL water. 1 µl of 2.5 µM RT primer were added and heated at 65°C for 5 min, cooled on ice and incubated in 20 µL RT mix (300 U Superscript III(Thermo Fisher Scientific, 18080044), 1X RT buffer, 500 µM dNTPs, 5 mM DTT) for 30 min at 55°C and heat inactivated at 70°C for 15 min. RT product was precipitated with ethanol as described above, resuspended in 10 µl water and resolved by 10% TBE-Urea gel. cDNA was isolated and eluted overnight in DNA elution buffer (DNA EB: 0.3 M NaCl, 1 mM EDTA), precipitated overnight, recovered and used for the 20 µL circularization reaction (100 U CircLigase™ ssDNA Ligase (Lucigen, CL4111K), 50 mM ATP, 2.5 mM MnCl_2_, 1X Circligase buffer) at 60°C for 2 hours. rRNA depletion was performed according to Ingolia et al.(Ingolia et al., 2012). Circularized, rRNA-depleted cDNA was used in the PCR using NEBNext® Ultra™ II Q5® Master Mix (NEB, M0544S) and NEBNext® Multiplex Oligos for Illumina® (NEB, E7335S). Half of the cDNA was used to determine the optimal PCR cycle number. The PCR products were resolved by 8% TBE gel (Thermo Fisher Scientific, Thermo Fisher Scientific). The other half of the cDNA was amplified by the determined PCR cycle, resolved by 8% TBE gel, and PCR products were cut, eluted in DNA EB, precipitated as described above, and recovered in 15 µl water. The quality and quantity of libraries were confirmed using DNA High Sensitivity Bioanalyzer (Agilent). Equal amounts of libraries were mixed and sequenced using NextSeq 550(Illumina). The quality control of the ribosome footprinting libraries is provided in the Figure S6.

To determine cytoplasmic RNA levels, total RNA was isolated from the cytoplasmic lysate (input that was loaded on the sucrose gradients for the abovementioned ribosome footprinting analysis) using Trizol LS according to the manufacturer’s instructions, except that precipitation with isopropanol was conducted overnight at -80°C. 1 µg RNA was used to make libraries with TruSeq Stranded Total RNA Library Prep Gold (Illumina, 20020598) according to the manufacturer’s instructions. The quality and quantity of libraries were confirmed using DNA High Sensitivity Bioanalyzer (Agilent) and sequenced using PE150, HiSeq 4000 sequencing platform at the Novogene Corporation Inc.

##### Alignment of sequencing reads

The 3′ adapters and low quality bases were trimmed by Cutadapt. Trimmed reads with length <15 nucleotides were excluded. The remaining reads were mapped to the mouse transcriptome using Bowtie (Langmead et al., 2009) with parameters: -a --best -m1 --strata. To construct the transcriptome, the annotation file from the ENSEMBL database (GRCm38) was used. For each gene, the mRNA with the longest CDS was selected. In the case of equal CDS length, the longest transcript was used. For read alignment, a maximum of two mismatches were permitted. To avoid ambiguity, reads that were mapped to multiple positions were excluded.

##### Aggregation plot of footprint reads near the start and stop codons

Firstly, the ribosome P-site was defined as positions +12, +13, and +14 from the 5’-end of the reads (the first position of the reads is recorded as 0). Then, for each mRNA, footprint reads in the CDS were counted. mRNAs with total reads in the CDS <32 or having empty codons (no observed reads) >90% were excluded. Ribosome densities at individual mRNA sites were normalized by the average density of the CDS. Then, ribosome densities with the same distance from the start or stop codon were averaged over the whole transcriptome.

##### Estimation of ribosome density on mRNAs

For each mRNA, a RPKM (Reads Per Kilobase of transcript, per Million mapped reads) value was calculated, and used to measure the relative ribosome density on individual mRNAs. mRNAs with RPKM values < 1 were excluded. For parallel pair-end RNA-seq, a FPKM (Fragments Per Kilobase of transcript per Million mapped reads) value was calculated. Ribo^ocp^ was calculated by dividing ribosome density by the corresponding mRNA level. mRNAs with fold change of ribosome density or mRNA level between samples higher than 2 (up-regulated) or lower than 0.5 (down-regulated) were defined as the groups of mRNAs with obvious difference in ribosome density or mRNA level.

##### Calculation of ribosome occupancy

For each mRNA, ribosome densities at individual codons of the CDS were normalized to the average density of the CDS. Second, ribosome occupancies at the same codons were averaged over the whole transcriptome. To estimate ribosome densities in different CDS regions, the CDS was divided into 100 bins, and ribosome occupancies at 61 codons (the three stop codons were not included) within the same bin were averaged over the whole transcriptome.

##### Calculation of pausing index

For each mRNA, a pausing index at the AUG codon was calculated. We defined the pausing index as the ratio of ribosome footprint reads at AUG codons over the other 60 codons. Upon the stress, mRNAs with pausing index increased up to 2-fold or higher were defined as paused mRNAs, and the remaining mRNAs were defined as the non-paused mRNAs. We defined the initiation pausing index as the ratio of ribosome footprint reads at the authentic initiation sites over the remaining of the CDS. Because majority of mRNAs showed dramatically increased initiation pausing (fold change of initiation pausing index > 2) upon the stress, the mRNAs with initiation pausing were defined as the mRNAs with fold change of initiation pausing index at the top 20%. As a control, the bottom 20% mRNAs were defined as the non-paused mRNAs.

##### Pathway analysis

For KEGG pathway analysis, the corresponding list of genes were fed into the STRING database (https://string-db.org/), and the pathways with the lowest FDR values were plotted. GO term analyses were performed with the GOEnrichment tool from Galaxy (Afgan et al., 2018) using a Benjamimi-Hochberg multiple test correction with a P-value cut off of 0.01 to generate a summarized list of enriched terms. PANTHER GO-Slim (Mi et al., 2019) was used as the gene ontology file along with the Mouse Genome Informatics GO annotation file, format 2.1, GO version 2020-03-23 (Ashburner et al., 2000; The Gene Ontology, 2019). Hierarchical clustering of q-values associated with each enriched GO term was performed using the NG-CMH Builder (Ward agglomeration, Euclidean distance) (Broom et al., 2017).

#### Polysome profile analysis

The lysate preparation and centrifugation conditions were identical to those described in the ribosome footprinting analysis. 12 fractions (1.2 mL) were collected, and RNA from each fraction was isolated using TRIzol LS reagent. An equal volume of RNA from each fraction was used for cDNA synthesis using the SuperScript™ III First-Strand Synthesis SuperMix for qRT-PCR(Thermo Fisher Scientific, 11752250). The relative quantity of specific mRNAs was measured by reverse transcriptase (RT)-quantitative polymerase chain reaction (qPCR) using VeriQuest SYBR Green qPCR Master Mix (Thermo Fisher Scientific, 756001000RXN) with the StepOnePlus Real-Time PCR System (Thermo Fisher Scientific, 4376599). For the list of primers see Table S1. For the studies of the migration of translation initiation factors, 1% formaldehyde was added to the lysis buffer and quenched with glycine (0.25 M) after 20 min on ice. Lysates were layered on 5%–20% sucrose gradients and centrifuged at 20,000 rpm for 20 h using an SW 32 Ti rotor. Twelve equal-size fractions were collected and precipitated with TCA (10%) at 4°C overnight. Precipitated proteins were washed twice with the wash buffer (volume 5:1 of acetone:1 M Tris-HCl at pH 7.4), dried, and dissolved in 1X SDS-PAGE sample buffer (50 mM Tris-HCl at pH 6.8, 5% β-mercaptoethanol, 2% SDS, 10% glycerol, 0.05% bromophenol blue). An equal volume of the dissolved proteins from each fraction was run on SDS-PAGE, transferred to membranes, and immunoblotted with specific antibodies.

#### Mitochondrial membrane potential measurement

MEF cells were cultured on coverslips overnight, followed by treatment with hyperosmotic stress and refreshed in isoosmolar media as indicated. After treatment, cells were incubated with 0.5 μM tetramethylrhodamine (TMRM; a cell permeant fluorescent red-orange dye as mitochondrial membrane potential indicator) (Thermo Fisher Scientific,T668) and Hoechst dye (Thermo Fisher Scientific, 62249) for 30 min at 37°C. After staining, cells were washed 3 times with warm PBS and fixed on coverslips with mounting reagent (DAKO,S3025). Cell images were taken by confocal microscopy (Fluoview FV100, Olympus) followed with ImageJ quantification of total TMRM signal and cell numbers from each image. Mitochondrial membrane potential from each image was calculated as the total TMRM signal normalized to the total cell number. At least 100 cells per image were counted in at least 4 fields per coverslip. Three replicates were analyzed for each indicated treatment.

#### Mitochondrial morphology and fragmentation measurement

MEF cells were cultured on coverslips overnight followed by treatment with hyperosmotic stress and recovery in isoosmolar media as indicated. After treatment, cells were fixed in 4% paraformaldehyde for 30 min and permeabilized with 0.1% Triton X-100. After blocking with 2% normal goat serum (Thermo Fisher Scientific, 10000C), fixed cells were incubated overnight at 4°C with anti-TOM20 primary antibodies (Proteintech,11802-1-AP) followed with 3 PBS washes. Secondary antibody with Alexa Fluor 488 (Thermo Fisher Scientific, A -11034) was applied to the coverslips followed by nuclei staining with Hoechst dye. After staining, cells were washed 3 times with warm PBS and mounted with mounting reagent. Cell images were taken by confocal microscopy (Olympus, Fluoview FV100) followed by ImageJ quantification of total cell numbers and cells with fragmented mitochondria from each image. At least 10 images were taken from each coverslip, and at least 100 cells per treatment were counted in the analysis. In this set of experiments, treatment of cells with 2 h of 10 μM of Carbonyl Cyanide Chlorophenylhydrazone (CCCP; Sigma-Aldrich, C2759) served as a positive control due to its known ability to severely induce mitochondrial fragmentation.

---

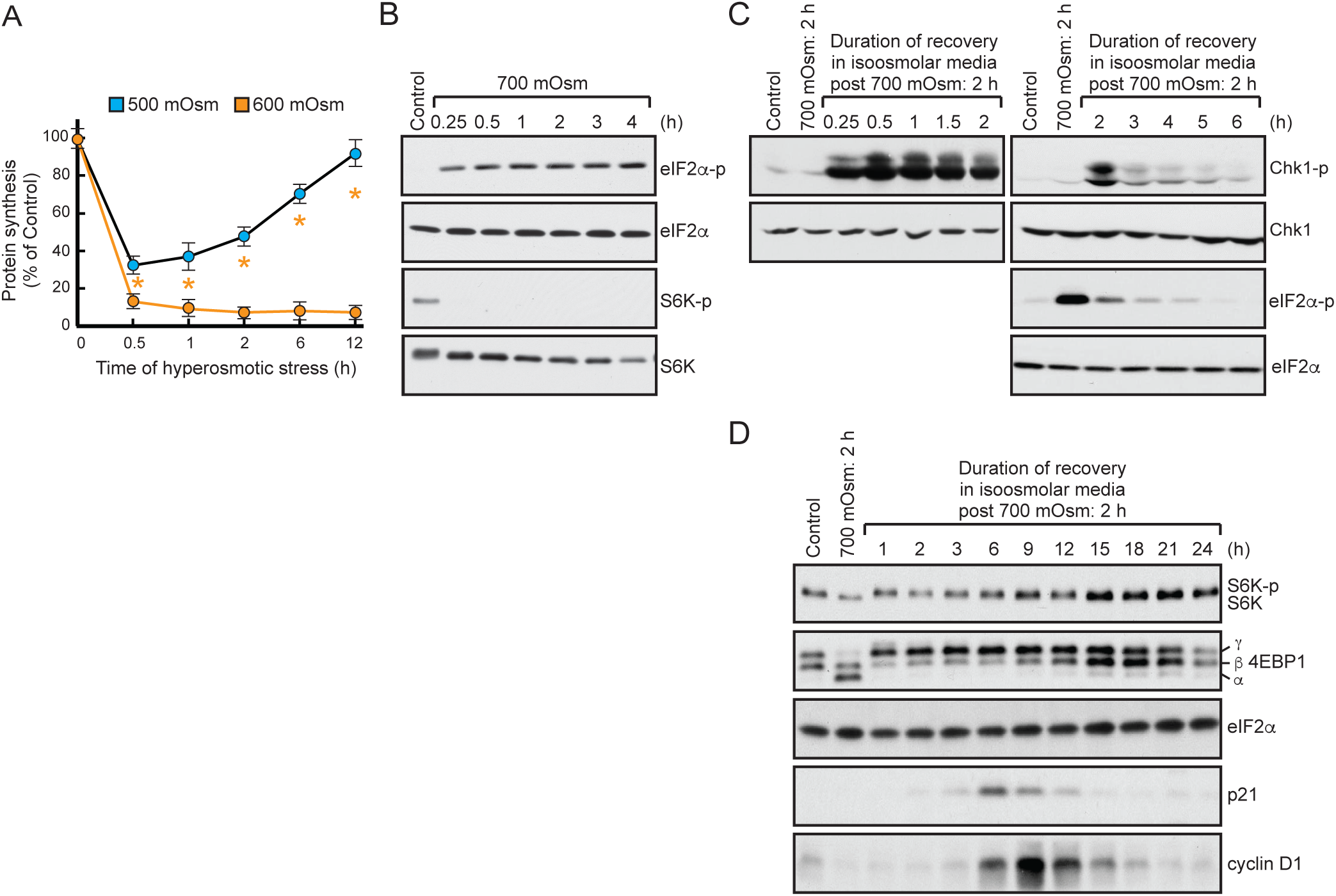
Figure S1, related to Figure 1

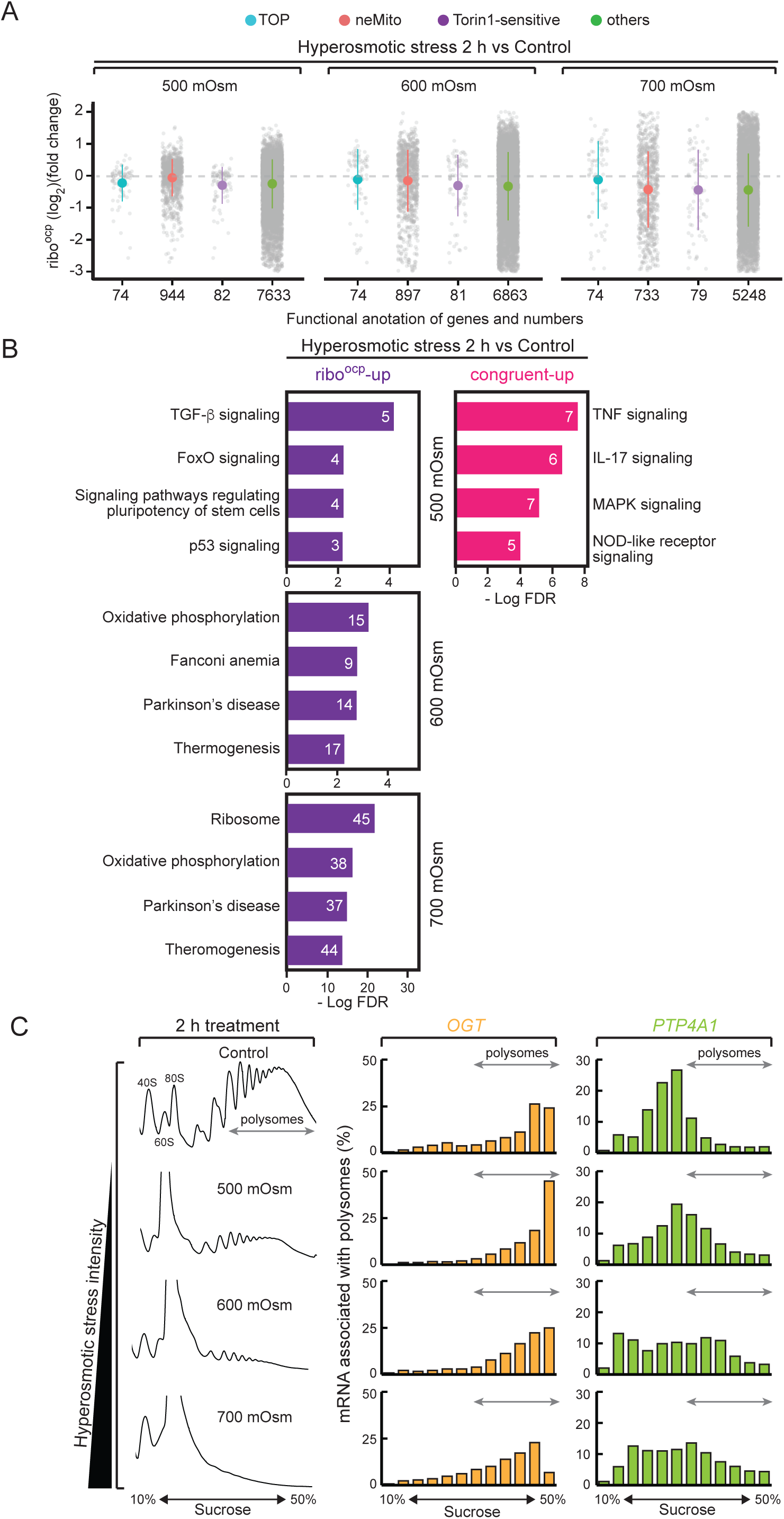
Figure S2, related to Figure 2

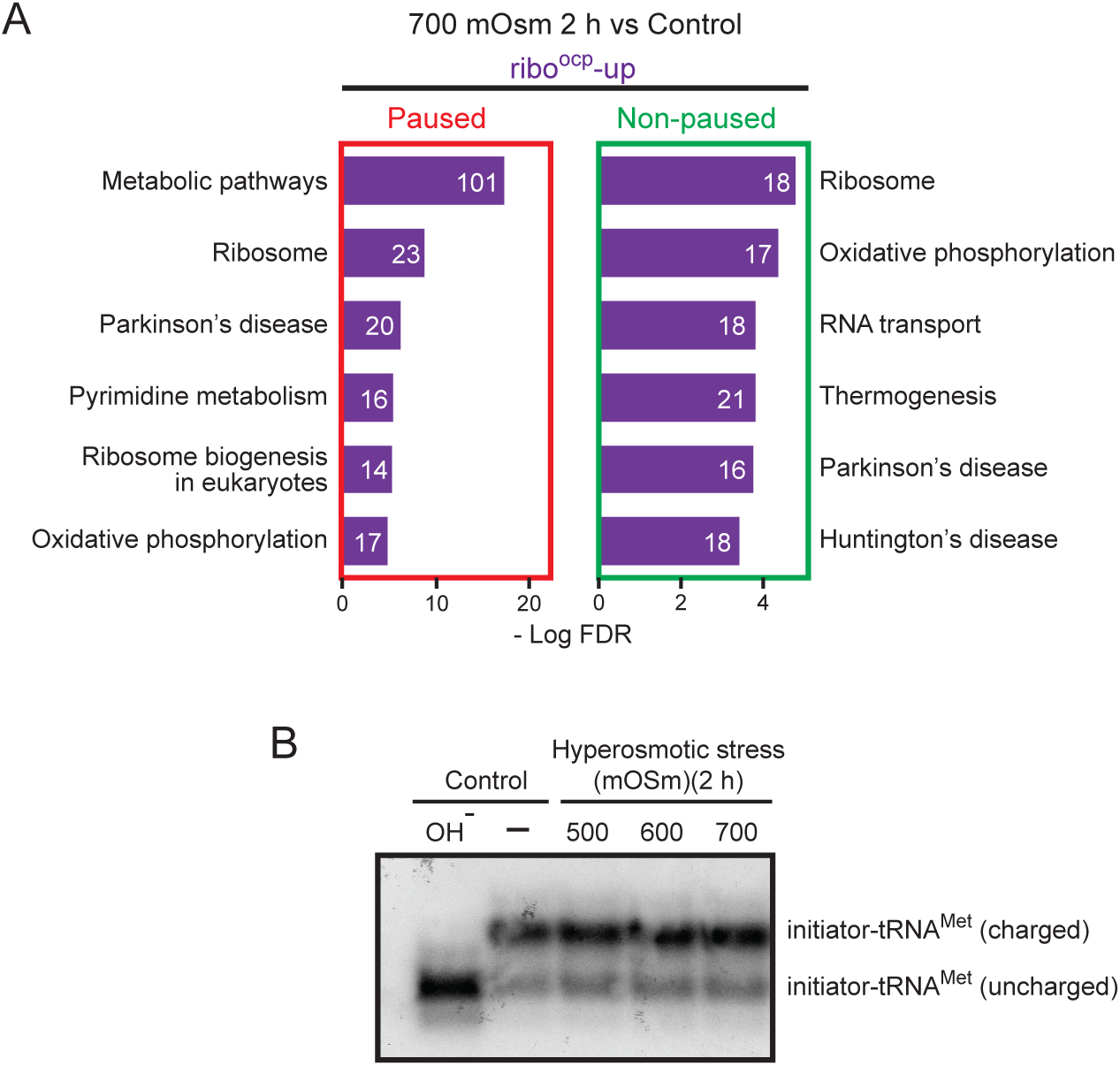
Figure S3, related to Figure 3

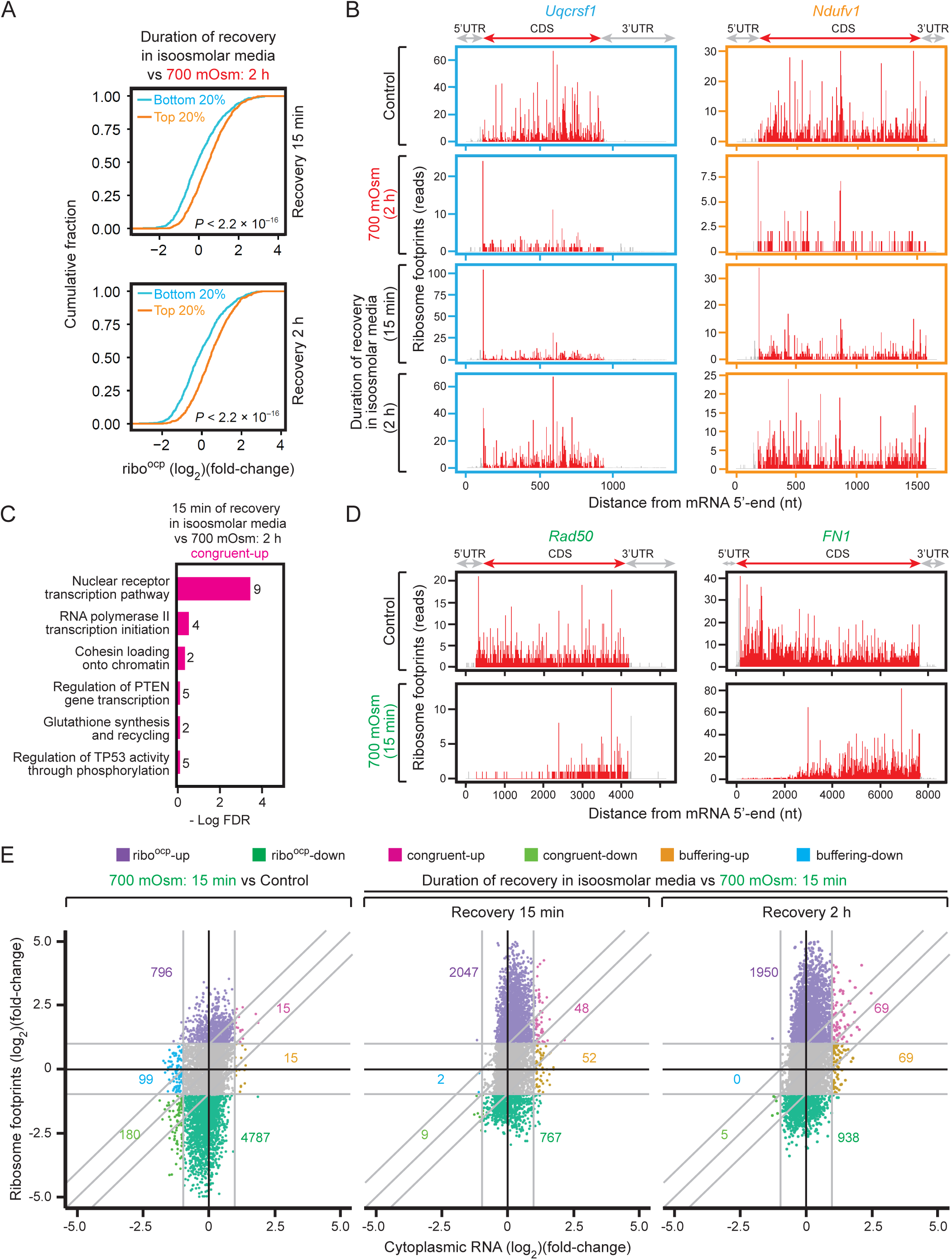
Figure S4, related to Figure 4

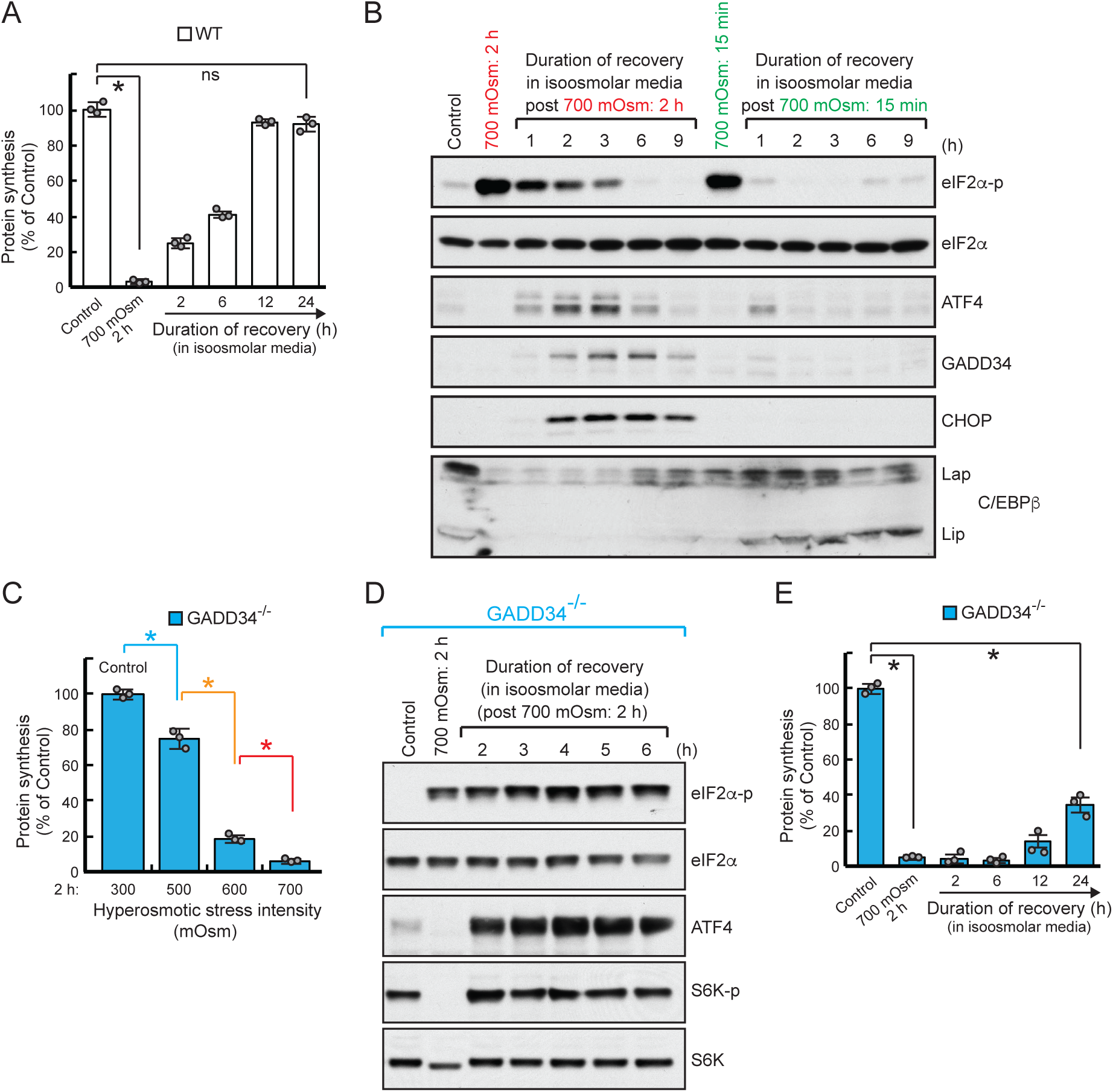
Figure S5, related to Figure 6

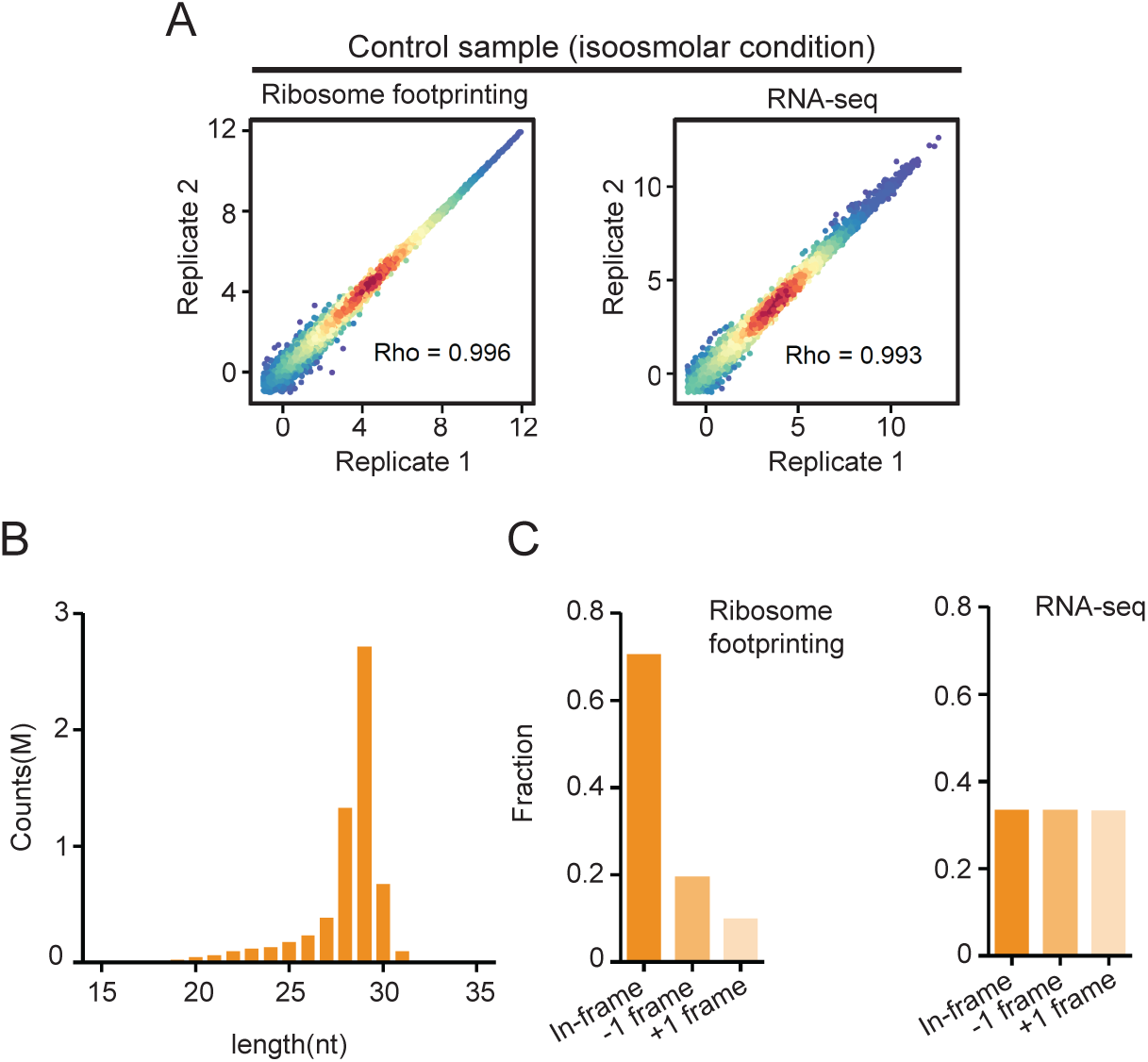
Figure S6, related to Materials and methods

